# Functional brain-wide network mapping during acute stress exposure in rats: Interaction between the lateral habenula and cortical, amygdalar, hypothalamic and monoaminergic regions

**DOI:** 10.1101/2022.05.10.491280

**Authors:** Laura Durieux, Karine Herbeaux, Christopher Borcuk, Cécile Hildenbrand, Virginie Andry, Yannick Goumon, Alexandra Barbelivien, Chantal Mathis, Demian Bataglia, Monique Majchrzak, Lucas Lecourtier

## Abstract

Upon stress exposure a broad network of structures comes into play in order to provide adequate responses and restore homeostasis. It has been known for decades that the main structures engaged during the stress response are the medial prefrontal cortex, the amygdala, the hippocampus, the hypothalamus, the monoaminergic systems (noradrenaline, dopamine, serotonin), and the periaqueductal gray. The lateral habenula (LHb) is an epithalamic structure directly connected to prefrontal cortical areas and to the amygdala, whereas it functionally interacts with the hippocampus. Also, it is a main modulator of monoaminergic systems. The LHb is activated upon exposure to basically all types of stressors, suggesting it is also involved in the stress response. However, it remains unknown if and how the LHb functionally interacts with the broad stress response network. In the current study we performed in rats a restraint stress procedure followed by immunohistochemical staining of the c-Fos protein throughout the brain. Using Graph Theory-based functional connectivity analyses, we confirm the principal hubs of the stress network (e.g. prefrontal cortex, amygdala, periventricular hypothalamus), and show that the LHb is engaged during stress exposure in close interaction with the medial prefrontal cortex, the lateral septum, and the medial habenula. In addition, we performed DREADD-induced LHb inactivation during the same restraint paradigm in order to explore its consequences on the stress response network. This last experiment gave contrasting results as the DREADD ligand alone, clozapine-N-oxide, was able to modify the network.

**GRAPHICAL ABSTRACT:** *GRAPHICAL ABSTRACT TEXT:* In this study, using immunohistochemical staining of the immediate early gene c-fos and graph theory-based functional correlational analyses, we aimed at unravelling the possible engagement of the lateral habenula (LHb) within the stress response network during acute stress exposure (10-min restraint) in rats. We found that the medial part of the LHb (LHbM) was preferentially engaged, and that this engagement was concomitant to this of structures such as the medial prefrontal cortex (mPFC), the insular cortex (Ins), hypothalamic (PVH) and thalamic (PVT) paraventricular nuclei, the extended amygdala, comprising the Bed nucleus of the stria terminalis (BNST) and the entire amygdala (AMG), as well as the dopaminergic ventral tegmental area (VTA) and the serotonergic dorsal raphe nucleus (RD). This suggests upon stressful situations the LHbM serves as a relay of cortical, thalamic, hypothalamic and temporal information, further transmitted to midbrain monoaminergic systems to probably initiate coping strategies. 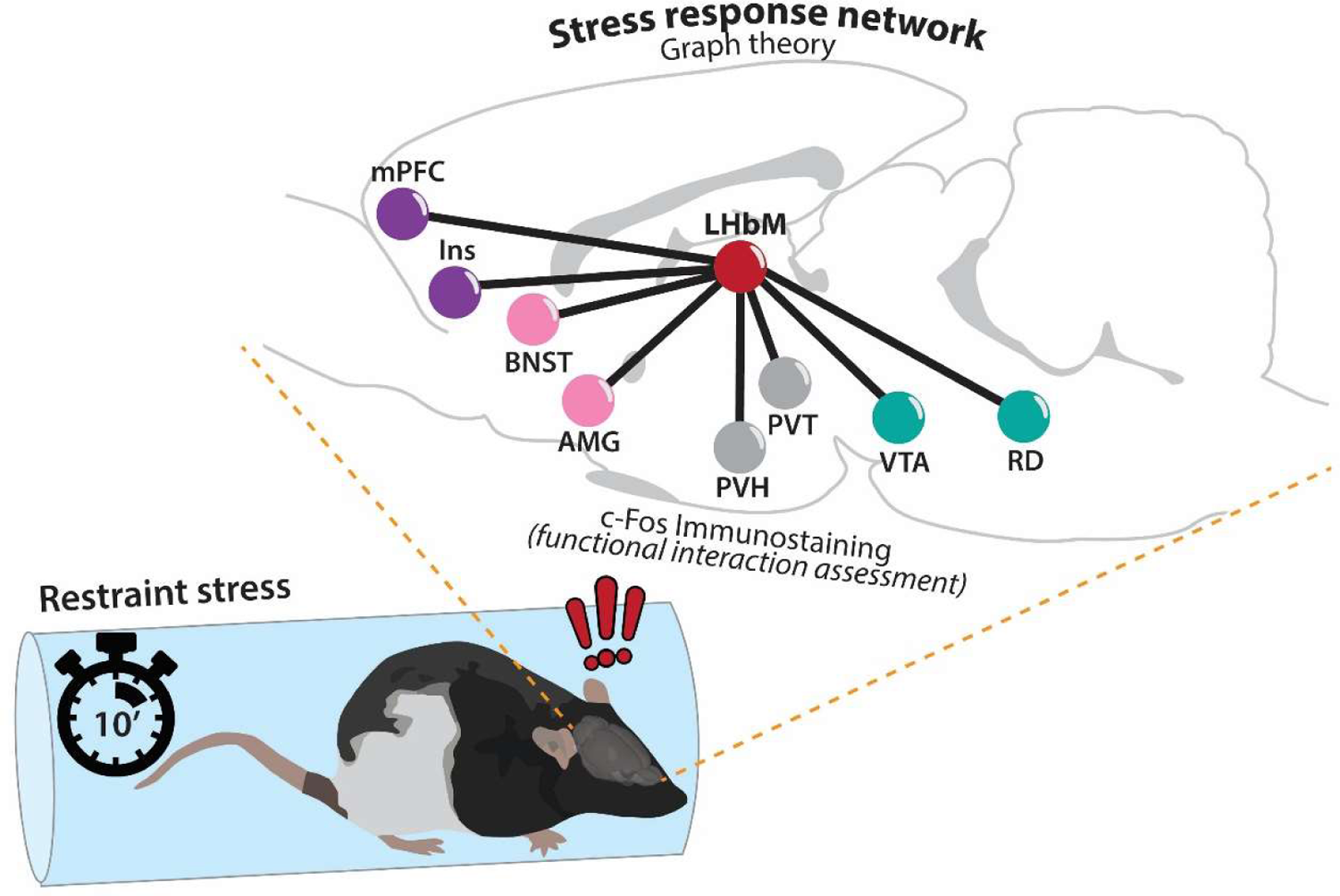

## INTRODUCTION

Adaptive response to stress exposure requires the coordinated activity of an extended brain network involved in cognitive, emotional, and motor processes (Joëls and Baram, 2009; Godoy et al., 2018). A large part of the studies addressing stress-induced changes in such functional network did so in the context of chronic stress, in order to better understand pathological conditions (e.g. Henckens et al., 2015; Magalhães et al., 2018). In humans, acute stressors engage key regions such as the insula (Ins), amygdala (AMG), anterior cingulate cortex (ACC), ventral striatum, prefrontal cortex (PFC), hippocampus (HPC), bed nucleus of the stria terminalis (BNST), thalamus, and the paraventricular nucleus of the hypothalamus (PVH) (Herman et al., 2003; Sousa, 2016). These structures – or their homologous regions - have also been shown to subserve physiological and behavioral responses to stress exposure in rodents (Herman et al., 2003; McEwen et al., 2015; Godoy et al., 2018).

In recent years, the lateral habenula (LHb), an epithalamic structure integrating forebrain information and modulating the activity of the main monoaminergic pathways, emerged as a prominent region for the control of the stress response (Hikosaka, 2010; Hu et al., 2020). The LHb is strongly activated by acute exposure to various stressors ranging from mild ones, like exposure to a novel environment (Wirtshafter et al., 1994; Durieux et al., 2020), to more severe such as electrical foot-shock, immobilization, tail pinch, and predator exposure (Chastrette et al., 1991; Wirtshafter et al., 1994; González-Pardo et al., 2012; Brown and Shepard, 2013; Dolzani et al., 2016; Zhang et al., 2016; Lecca et al., 2017; Barrett and Gonzalez-Lima, 2018; Durieux et al., 2020). Interestingly, in rodents the LHb directly or indirectly interacts with the above-cited key regions of the stress response. Regions which send direct projections onto the LHb comprise the entire PFC (Kim and Lee, 2012), and the extended amygdala, including the BNST (Dong and Swanson, 2004, 2006) and central amygdalar nucleus (Zhou et al., 2019). The LHb and HPC, although they do not share direct connections, likely communicate and functionally interact (Aizawa et al., 2013; Goutagny et al., 2013; Baker et al., 2019). The LHb has been shown to modulate the hypothalamus-pituitary-adrenal (HPA) axis (one of the primary effectors of the stress response; Selye, 1950; de Kloet et al., 1998), and therefore most likely the activity of the PVH (Murphy et al., 1996; Jacinto et al., 2016; Mathis et al., 2018). In addition, the LHb, as shown in rodents, cats and monkeys, can modulate the main neurotransmitter systems involved in the stress response, *i.e.,* dopamine (Christoph et al., 1986; Ji and Shepard, 2007; Lecourtier et al., 2008), noradrenaline (Kalen, 1989a; Cenci et al., 1992), and serotonin (Reisine et al., 1982; Kalen et al., 1989b). The link with the serotonergic system is of particular interest as inhibition of the LHb-dorsal raphe nucleus pathway in rats reduces passive stress response (Coffey et al., 2020). The LHb also interacts with other regions to process emotional responses. For example, in mice inhibition of the lateral hypothalamus-LHb pathway alters escape behavior upon unpredictable exposure to foot-shocks (Lecca et al., 2017), and stimulation of the pathway connecting the entopeduncular nucleus to the LHb (Shabel et al., 2012), the LHb to the ventral tegmental area (Lammel et al., 2012), and the LHb to the rostromedial tegmental nucleus (Stamatakis and Stuber, 2012), induces aversive responses. Finally, probably through an indirect activation of rostral medullary raphe neurons, the LHb has been involved in rats in the control of stress-induced hyperthermia, a component of the physiological stress response (Ootsuka et al., 2017).

To date no study has been devoted to unravelling if and how the LHb is part of the broad network involved in the stress response. In the current study we used acute exposure to restraint stress as a behavioural paradigm and specifically aimed to describe: 1) the network activated by restraint and the position of the LHb within this network, and 2) the consequences of DREADD-induced LHb inactivation on the connectivity of the network. To this aim we quantified the level of expression of the c-Fos protein in a large number of brain regions, including the key structures involved in the acute stress response. The immediate-early gene c-Fos appeared relevant to explore as it is the best characterized and most widely used tool for functional anatomical mapping in rodents, given its rapid activation by various stimuli (Kovács, 1998; Hudson, 2018). The quantification of the density of c-Fos positive (c-Fos+) cells throughout the brain, and the exploration of between-structure correlations can reveal functional interactions between structures belonging to a given network, as several studies previously argued (e.g., in fear memory; Wheeler et al., 2013; Vetere et al., 2017). This level of analysis can be achieved applying a Graph theory approach. Graph theory is a mathematical field allowing to analyze complex networks (Bullmore and Sporns, 2009). The main principle of this approach is to consider the structures as nodes of a given network, nodes that are connected through edges (the functional variable, in our study, c-Fos+ cells density correlations). Then, different parameters of the network can be revealed, such as the principal hubs of the network, though strength analysis, or the presence of sub-networks, which could be engaged in different aspects of the ongoing process, through the extraction of modules/communities (see Bullmore and Sporns 2009 for examples). Finally, in animals exposed to restraint, we assessed the level of plasmatic corticosterone, before and following restraint, in order to control for the effectiveness of such procedure and to further investigate if the LHb modulates the HPA axis upon stress exposure.

## MATERIALS & METHODS

### Animals

This experiment, authorized by the French authorities (APAFIS#7114), was carried out with 44 five months old male Long–Evans rats (Janvier Labs, France) issued from a previous study (Durieux et al., 2020). They were housed in pairs on a 12 h light/dark cycle (lights on at 7:00 A.M.) with *ad libitum* access to food and water. Rats were singly housed five days before the start of the experiment and, the last three days, were familiarized to the holding procedure later used to collect blood samples. The experiment took place between 9:00 and 12:00 PM. A schematic representation of the experimental procedure is provided in **Figure 1**.

**FIGURE 1.**
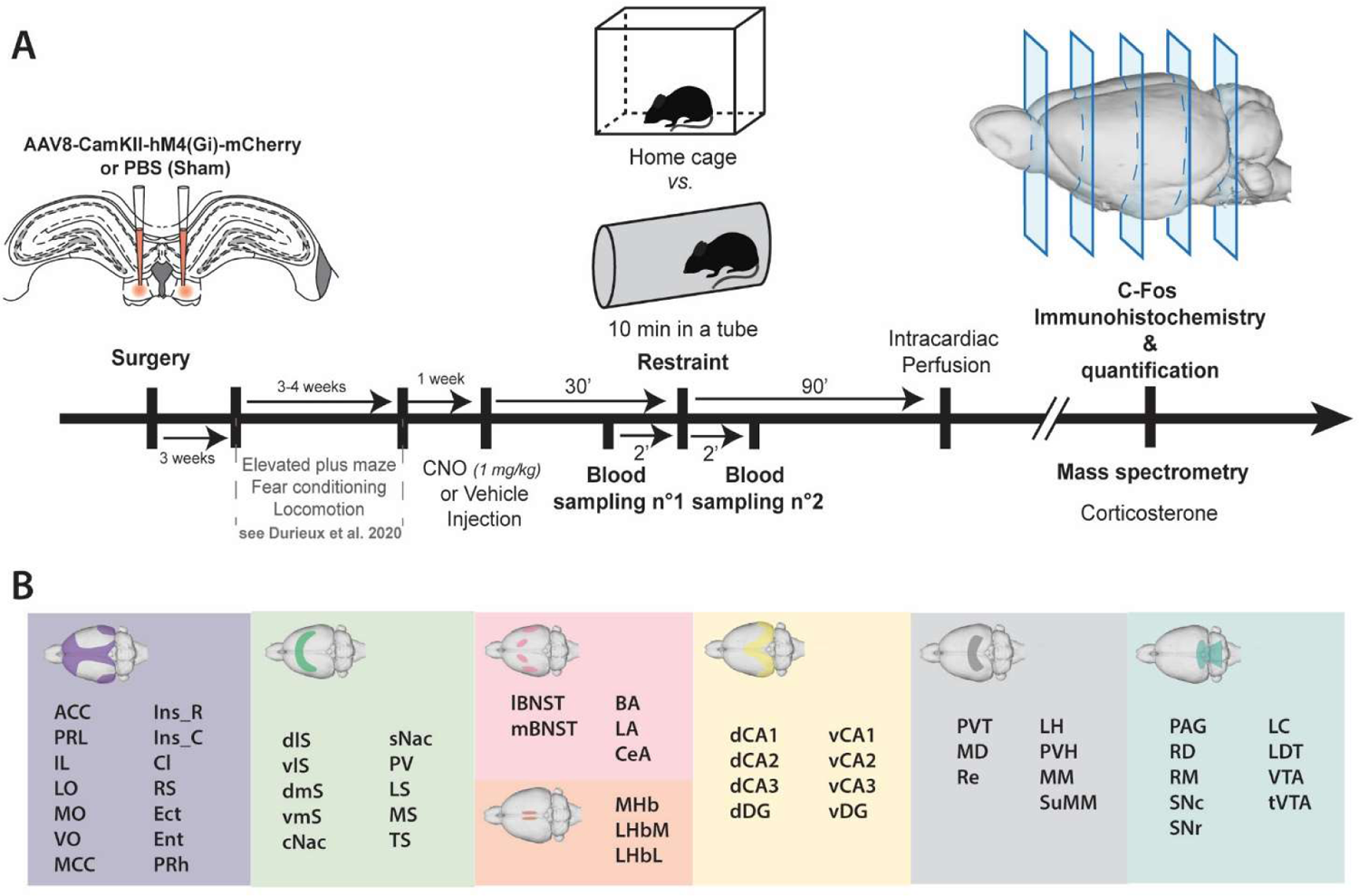
(**A**) Timeline of the experiment. Rats of the present study are those of a previously published study (Durieux et al. 2020). They underwent surgery 7 to 9 weeks prior to the present experiment, and then went on several tests including elevated plus maze, fear conditioning and locomotor activity assessment. One week elapsed between the end of the previous set of experiment and the restraint procedure during which they were kept in a tube for 10 min. Controls remained in their home cage. The injection of CNO (1 mg/kg) or vehicle was performed 30 min before. In restrained animals, blood samples were collected once 2 min before and once 2 min after restraint. Eighty min following restraint, animals were deeply anesthetized, intracardiac perfusion was performed and brains were removed. Blood samples were run though a mass spectrometer to assess corticosterone levels. Brains were cut, and immunolabeling of the c-Fos protein was performed on 40 µm-thick floating sections. The density of c-Fos+ cells was calculated in each region of interest. (**B**) Regions of interest comprised cortical (purple), basal ganglia and septal (green), extended amygdalar (pink), habenular (orange), hippocampal (yellow), thalamic and hypothalamic (grey), and brainstem (blue) regions.

### Surgery

Surgery was conducted 7-9 weeks before the experiment as described in detail in Durieux et al. (2020). Shortly, under isoflurane anesthesia [4% for induction; 1.5 % throughout surgery] and painkiller supply (meloxicam, 1 mg/kg, *s.c.*), rats were bilaterally microinjected with either the viral vector (AAV8–CamKII–hM4(Gi)–mCherry; Viral Production Unit, Universitat Autonoma de Barcelona; hM4 animals, n=22), or phosphate–buffered saline (PBS animals; n=22) at the two following coordinates and volumes: 1) anteroposterior (AP) = – 3.3 mm from Bregma, mediolateral (ML) = ± 0.7 mm from the midline of the sagittal sinus, dorsoventral (DV) = – 4.5 mm from dura (0.2 µL); 2) AP = – 3.5 mm from Bregma, ML = ± 0.7 mm from the midline of the sagittal sinus, DV = –4.4 mm from dura (0.15 µL).

### Drug treatments

Rats were allocated to CNO (1 mg/kg; dissolved in 0.9% NaCl–0.5% DMSO; *i.p.*) or vehicle (Veh, 0.9% NaCl–0.5% DMSO, *i.p.*) treatment as in our previous study (Durieux et al., 2020; see Supporting information Table S1). Thirty minutes after drug injection, animals of each treatment condition were either exposed to blood sampling and restraint [restraint stress (RS) groups], or remained undisturbed in their home cage [home cage (HC) group]. More precisely, sixteen hM4 animals and sixteen PBS animals were administered CNO and pseudo-randomly distributed in the two behavioural conditions as follows: home-cage condition [HC; hM4–CNO-HC group (n=8) and PBS–CNO-HC group (n=8), respectively] or restraint stress condition [RS; hM4–CNO-RS group (n=9) and PBS–CNO-RS group (n=7), respectively]. Six rats injected with PBS and six rats injected with AAV8–CamKII–hM4(Gi)–mCherry (to control for the potential effects of the expression of both the hM4 receptors and the mCherry fluorophore) were administered vehicle (Veh); those rats composed the two Veh control groups, such as it was done in Durieux et al. (2020), each composed of 3 rats injected with PBS and 3 rats injected with AAV8–CamKII–hM4(Gi)–mCherry, and named (PBS or hM4)-Veh-HC and (PBS or hM4)-Veh-RS according to the behavioral condition they were pseudo-randomly assigned to, home cage and restraint stress respectively.

### Restraint

Rats were gently placed in an opaque grey PVC tube (length, 80 cm; diameter, 13.5 cm) including ventilation holes at each end. They remained in it for a duration of 10 min.

### Plasmatic CORT concentration assessment

#### Blood sampling

Blood sampling was performed in RS groups 2 min before (pre-restraint) and 2 min after (post-restraint) the restraint procedure. Rats were put on a table with their head positioned in a folded towel, and gently held in place. The caudal vein was lightly incised with a razor blade and blood, approximately 250 µl each time, was collected through a heparinized capillary tube (Microvette CB 300) while gently stroking the tail from the base to the tip. The pre-restraint incision was performed about 3 cm from the tip of the tail, and the post-restraint one was performed 2 cm higher. As soon as blood was collected, the tubes were placed in ice (4°C) before being centrifuged (3000 rpm at 4°C during 4 min) to collect approximately 150 µl of plasma, which was then stored at −80°C until analysis was performed.

#### CORT preparation

For each sample, 50 µl of plasma were added to 10 µl of D4-CORT (50 pmole/10 µl, Sigma-Aldrich, St. Quentin Fallavie, France) in 99.1% H2O / 0.1 % formic acid (v/v). Then, proteins were precipitated by adding 1 ml of acetonitrile (ACN 100 %). Two successive centrifugations were performed at 20,000 g (4°C, 20 min) and, the resulting supernatant was recovered and dried under vacuum (SpeedVac, Thermo Fisher). Samples were re-suspended in 30 µl of 20 % ACN / 0.1 % of formic acid, followed by a last centrifugation (20,000 g, 10 min). Supernatants were recovered and kept at −80°C until LC-MS/MS analysis.

#### Liquid Chromatography Tandem-mass Spectrometry (LC-MS/MS)

Analyses were performed with a Dionex Ultimate 3000 HPLC system (Thermo Electron, Villebon-sur-Yvette, France) coupled with a triple quadrupole Endura mass spectrometer (Thermo Electron). Samples were loaded onto a ZORBAX SB-C18 column (150 x 1 mm, 3.5 μm, flow of 90 µl/min; Agilent, Les Ulis, France) heated at 40°C. LC and MS conditions used are detailed in Supporting information **Table S2-3**. Qualification and quantification were performed using the multiple reaction monitoring mode (MRM) according to the isotopic dilution method (Ho et al., 1990). Identifications of the compounds were based on precursor ions, selective fragment ions and retention times obtained for the heavy counterpart (*i.e.,* IS). Selection of the monitored transitions and optimisation of collision energy and RF Lens parameters were determined automatically (Supporting information **Table S3**). Xcalibur v4.0 software was used to control the system (Thermo Electron).

### Brain tissue preparation and section processing

Eighty minutes after the end of the restraint stress procedure, rats were deeply anesthetized with a pentobarbital overdose (120 mg/kg, *i.p*.). Following intracardiac perfusion of phosphate-buffered saline (PBS, 0.1M) and then 4% paraformaldehyde (PFA)-PBS solution (pH 7.4; 4°C), brains were removed, post-fixed in 4 % PFA-PBS (4°C, 2 h), transferred into a 0.1 M PBS–20 % sucrose solution (4°C, 48 h) and subsequently frozen (isopentane, −40°C, 1 min). Serial 40 μm-thick free–floating sections were cut – 1 every 120 µm – in the coronal plane at – 20°C and stored in a cryoprotectant solution at – 20°C (see Supporting information **Table S4** for a detailed description of the coordinates of the boundaries of the different structures investigated according to the Paxinos and Watson stereotaxic atlas). In animals injected with the DREADD viral solution, within the block containing the LHb, one every three sections were directly collected on gelatin-coated slides in order to assess the extent of mCherry expression.

### c-fos Immunohistochemistry

The immunohistochemistry protocol used was performed such as during our previous study (see Durieux et al., 2020 for details). In summary, c-Fos proteins were bound by primary anti–Fos rabbit polyclonal antibody (1:750, polyclonal rabbit antibodies; SYSY; ref: 226 003, Synaptic System). The secondary antibody was a biotinylated goat anti-rabbit antibody (1:500, Biotin-SP-conjugated affiniPure Goat anti-rabbit IgG, ref: BA1000, Vector). Staining was performed with the avidin–biotin peroxidase method (Vectastain ABC kit, PK 6100; Vector Laboratories, Burlingame, CA, USA). Sections were subsequently mounted on gelatin-coated slides, dehydrated by incrementally concentrated alcohol baths (70%, 90%, 95%, 100%, 100%), covered with Clearify (Americain MasterTech Scientific) and covered with Diamount (Diapath S.P.A). In addition, in rats in which LHb contained hM4(Gi) receptors and which were administer CNO during HC and RS conditions, we performed fluorescent c-fos immunohistochemistry staining on slices containing the LHb. Slices (from 3 to 6 per rat) adjacent to the ones used for the previously performed DAB/bright field c-Fos immunohistochemistry were chosen. Floating slices were exposed to a primary anti-Fos guinea pig antibody (1:1000, ref: 226005, Synaptic Systems) followed by exposure to a goat anti-guinea pig secondary antibody (1:500, ref A-11073, ThermoFisher) coupled to Alexa Fluor 488. Slices were then mounted on gelatin-coated slides and covered with a DAPI–fluoromount medium. Sections were scanned and saved in NDPI format using a NanoZoomer S60 (Hamamatsu) at x 20 thanks to the associated software (NDP.scan, Hamamatsu) [c-Fos, 488 nm (FITC filter); mCherry, 594 nm (TRITC filter)].

### Quantification

Quantification of c–Fos positive (c-Fos+) cells revealed with avidin-biotin method was performed in 56 structures (see Supporting information **Figure S1-2**) using a semi–automated method with ImageJ (Free License, Wayne Rasband, Research Services Branch, National Institute of Mental Health, Bethesda, Maryland, USA; testing of the accuracy of the method is presented in Supporting information **Figure S3**) such as described in Durieux et al. (2020). We took from 3 to 20 sections per structure depending on their rostro-caudal extent. Both hemispheres were pooled. Results were expressed as mean number of c-Fos+ cells by mm² (for more details see Supplementary information). Quantification of LHb cells that were c–Fos positive (c-Fos+) and mCherry positive (mCherry+) was performed using the software QuPath (Bankhead et al., 2017). First, the LHb was manually outlined on each slices (3 to 6 per rat) using outlines identical to those used for the previously performed DAB/bright field c-Fos immunohistochemistry. Then we extracted the number of fluorescent cells in the two filters based on the detection cells algorithm; the settings used were as follows: background radius (8 µm), sigma (1.5 µm); the cell area was expected to be between 10 and 400 µm so that the threshold was set to 10. Once the detection of the cells terminated the area of the overlay was saved along with the number of detected cells in each wavelength. The density of both c-Fos+ and mCherry+ cells (in number/mm²) was calculated dividing the number of the cells detected by the area delimited by the overlay. The number of cells that were both c-Fos+ and mCherry+ (colocalization; c-Fos+/mCherry+) was counted manually in the same defined areas.

### Statistical analyses

Between-group [(PBS or hM4)-Veh, PBS-CNO, and hM4-CNO in the RS condition] comparison of plasmatic corticosterone (CORT) levels was performed using a two-way ANOVA with Time (pre-vs post-restraint) as the repeated measure. For each brain region, c-Fos raw data (c-Fos+ cells densities) were analyzed with ANOVAs with Group [(PBS or hM4)-Veh, PBS-CNO, hM4-CNO] and Condition (HC and RS) as between-subject factors. Post hoc Newman-Keuls multiple range test was used when appropriate. In all analyses significance was set for *p* < 0.05.

### Functional connectivity analysis

We used this analysis method to model the functional network during stress response. For this purpose, only c-Fos data of the RS groups were further analyzed using network modeling [(PBS or hM4)-Veh-RS, n=6; PBS-CNO-RS, n=7; hM4-CNO-RS, n=7]. They were processed using Python (version 3.7.5, PyCharm edition community – free; https://www.jetbrains.com/fr-fr/pycharm/) with helps from several toolbox. For general coding, we used pandas (manipulating data as DataFrame and XLSX/CSV export/import; https://pandas.pydata.org/), numpy (managing matrix; https://numpy.org/), Scipy (for general statistic purposes; https://www.scipy.org/). The others Python toolbox used during the analysis for precise goal are cited when necessary. Correlations were calculated using Korr (https://pypi.org/project/korr/) according to the Pearson method. The correlation matrix was plot using matplotlib (https://matplotlib.org/) and seaborn (https://seaborn.pydata.org/). The 3D Rats brains representations used have been created from The Scalable Brain Atlas (Bakker et al., 2015).

The network modeling was processed using NetworkX toolbox (https://networkx.org/). The graphs were undirected and unsigned (absolute correlations were used as weights). All analyses were done on full weighted networks. Graph theoretical metrics, *i.e.,* the strength of each node in the network, the modularity (based on Louvain algorithm module/community detection, see below), the average clustering coefficient, and the average of edge weights, were computed using NetworkX. We used classical bootstrap analysis to estimate the variance of the population in groups or in random model. The random model was computed by shuffling the row and the column of the correlation matrix keeping the main diagonal intact (so that each structure remains the same in the random model). Between-group comparisons were done using the permutation test. The permutation test is a statistical test under the null hypothesis based on the calculation of the shuffling of the two compared groups (creating the null hypothesis) and comparing the given distribution of the p value to the *p* value of both original groups. Only data showing a significance difference (*p* < 0.05) in both tests (bootstrap and permutation) were estimate significantly different.

The allegiance matrices were computed using the Louvain method which allows to unfold communities in large networks (Python package https://pypi.org/project/python-louvain/ called by NetworkX). The modules (or communities) were calculated for each bootstrap, modulating the resolution of Louvain algorithm from 0.88 to 1 over 10 repetitions, so that the number of communities varied from 2 to 5 depending on the group considered (see Supporting information **Figure S4**). The allegiance represents the probability that two structures are in the same community across bootstrap iterations over all possible pair of structures. The allegiance matrix also displays modules, allowing us to extract the communities on this matrix and study which structures may be activated in a functionally coordinated manner. This calculation can highlight high synchronicity and high functional connectivity between structures of the same community. Once the allegiance matrix calculated, the allegiance communities were extracted using the Louvain algorithm (resolution of 1). The allegiance communities of the (PBS or hM4)-Veh-RS group was extracted and the “within-module strength Z-score” evaluated (see Supporting information **Figure S4**).

Between-group comparisons were based on the communities extracted from the (PBS or hM4)-Veh-RS group. The average strength of the structures of each community was calculated for each group and compared (PBS or hM4)-Veh-RS vs PBS-CNO-RS, (PBS or hM4)-Veh-RS vs hM4-CNO-RS and PBS-CNO-RS vs hM4-CNO-RS) using the non-parametric test, Wilcoxon-Mann-Whitney. The differences were considered significant for p<0.05. On the same principle, we also tested the possible differences between the PBS-CNO and the hM4-CNO communities based on PBS-CNO communities. For each group we assessed the distribution of the structures of interest in the different communities, using the so-called confusion matrices, based on the principle that communities do not have a hierarchical level, so that one may wonder which community of a given group best represents the community of another group. We calculated the Jaccard index gauging the similarity and the diversity of a sample set.

## RESULTS

### Histology

Following histological verification (extend of mCherry staining), groups were composed as follows: (PBS or hM4)-Veh-HC, n = 6; (PBS or hM4)-Veh-RS, n = 6; PBS-CNO-HC, n = 8; PBS-CNO-RS, n = 7; hM4-CNO-HC, n = 4; hM4-CNO-RS, n = 7. The extent of the expression of mCherry in hM4-CNO animals is represented in Supporting Information **Figure S5**.

### Blood CORT release

The two-way ANOVA indicated no significant effect of Group (*F*_2,17_ = 0.13; *p* > 0.8), a significant effect of Time (*F*_1,17_ = 235.76; *p* < 0.0001), and no interaction between the two factors (*F*_2,17_ = 0.21; *p* > 0.8) (**Figure 2**). This indicates that whereas the restraint procedure was indeed stressful, leading to a marked increase of CORT release, LHb inactivation had no impact on such response, although a ceiling effect remains a possibility.

**FIGURE 2.**
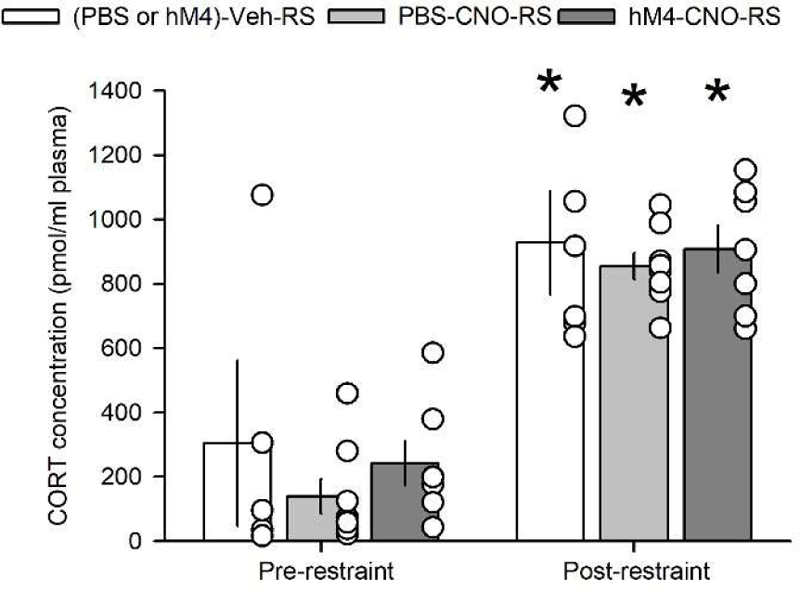
Effect of restraint and LHb inactivation on CORT plasmatic concentration. Mean CORT plasmatic concentration (± S.E.M) before (left) and after (right) restraint in (PBS or hM4)-Veh-RS (white bars), PBS-CNO-RS (light grey bars) and hM4-CNO-RS (dark grey bars) groups (white circles represent individual values). <u>Statistics</u>: \**p* < 0.05 vs. pre-restraint in the same group.

### c-Fos+ cell densities

c-Fos expression was high in almost all structures in restraint groups but also in the HC hM4-CNO group (**Figure 3A**; see also Supporting Information **Table S5** for the raw data and **Tables S6** and **S7** for statistics).

**FIGURE 3.**
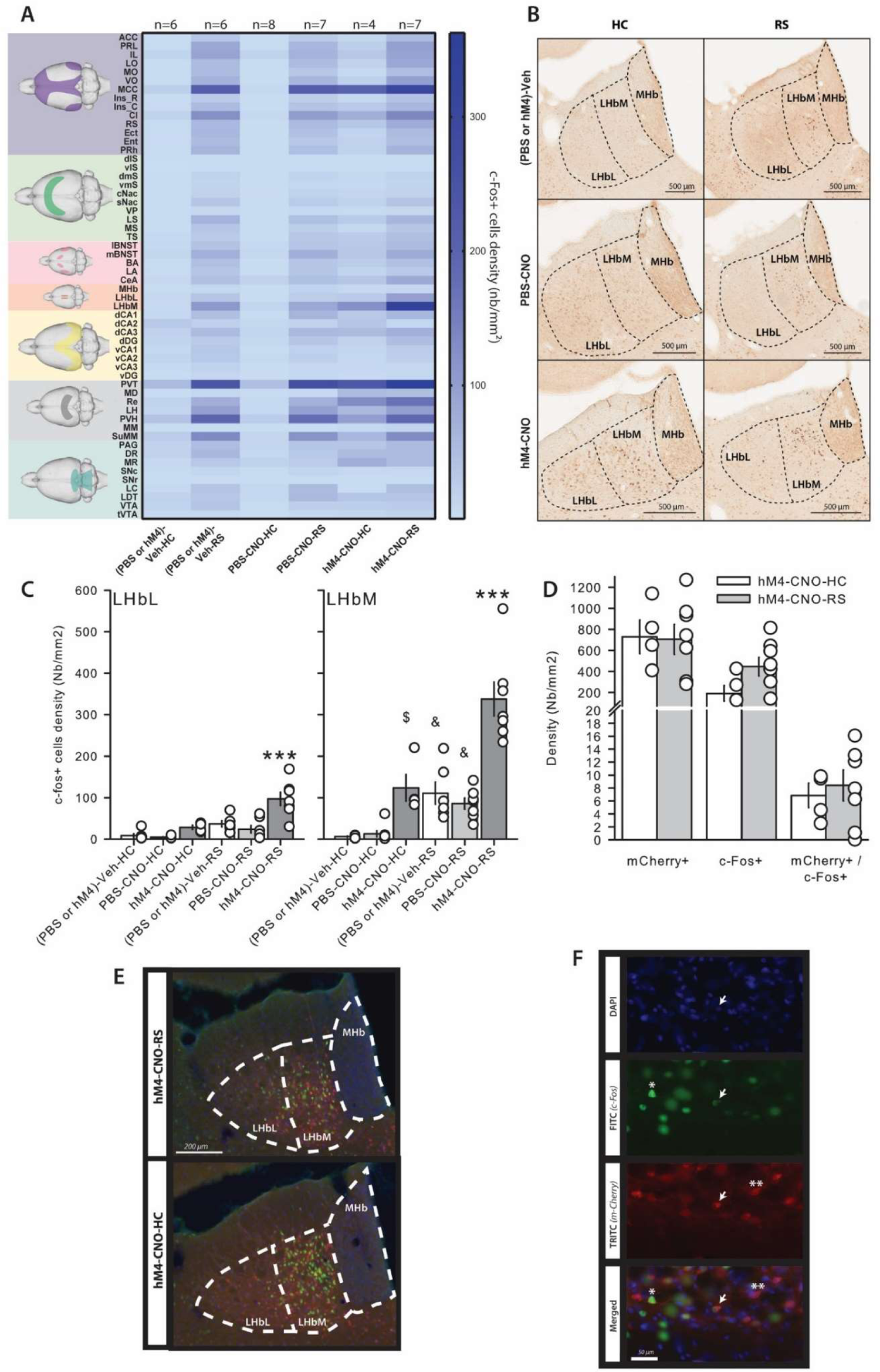
Effects of restraint and LHb inactivation on c-Fos expression. (**A**) Heatmap of mean c-Fos+ density per structures for all groups. (**B**) Photography of c-Fos labeling in the entire habenula in (PBS or hM4)-Veh (top), PBS-CNO (middle), and hM4-CNO (bottom) groups in the HC (left) and RS (right) conditions. (**C**) Bar plot of c-Fos+ density (mean ± S.E.M) in the LHbL and LHbM (white circles represent individual values). (**D**) Density (mean ± S.E.M) of mCherry+, c-Fos+, and mChery+/c-Fos+ doubly labeled neurons within the LHb of animals injected with the viral vector AAV8–CamKII–hM4(Gi)–mCherry and injected with CNO either in the HC (white bars) or in the RS (light grey bars) condition (white circles represent individual values). (**E**) Representative image of the c-Fos+ (green) and mCherry+ (red) neuronal population in the LHb of a hM4-CNO-RS (top) and a hM4-CNO-HC (bottom) rat. (**F**) Magnification of LHb neurons showing the different neuronal population such as they were observed following c-Fos immunostaining, i.e. a c-Fos+ neuron (*), a mCherry+ neuron (**), and a doubly labelled mCherry+/c-Fos+ neuron (pointed to by a white arrow). <u>Statistics</u>: \*\*\**p* < 0.001 vs. all other groups; ^&^*p* < 0.05 vs. the corresponding HC group; ^$^*p* < 0.05 vs. the other HC groups (Newman-Keuls post-hoc test following ANOVA).

The restraint-induced increase in c-Fos expression was evidenced by significant Condition effect in almost all structures at the exception of the dlS, vlS, dCA2, MD, MM, SNc, SNr, and LDT; in the latter, a significant Group effect due to higher c-Fos+ cells density in the hM4-CNO group than in both control groups was observed in the dlS, vlS, MD, and MM, and a significant interaction without main effects of each factor due to a higher c-Fos+ cells density in the hM4-CNO-RS group than in the PBS-CNO group of the same condition (*p* < 0.05) in dCA2. In regions showing activation upon restraint, the analyses showed a significant Group effect - also due to higher c-Fos+ cells density in hM4-CNO group compared to the two control groups which did not differ-in the PRL, LO, VO, MCC, Ins_R, Ins_C, Cl, RS, vmS, PV, LS, TS, BA, CeA, dCA3, PVT, Re, and tVTA; in the LH the ANOVA also indicated a significant interaction due to higher c-Fos expression in hM4-CNO-HC group as compared to both HC control groups (*p* < 0.001 for each comparison). The lack of difference between the (PBS or hM4)-Veh and PBS-CNO groups indicates that the effects observed in the hM4-CNO group are specific to the action of CNO on hM4(Gi) receptors. However, in the LS, the Group effect was also due to lower c-Fos+ cells density in PBS-CNO and, in the RD, it was only due to lower c-Fos+ cells density in PBS-CNO group than the others, suggesting that CNO was not devoid of effects in our conditions albeit in only a few regions. In both LHb subdivisions (**Figure 3B-C**), the ANOVA indicated significant effects of Condition (LHbL: *F*_1,3_2 = 25.44, *p* < 0.00001; LHbM: *F*_1,32_ = 43.003, *p* < 0.00001) and Group (LHbL: *F*_2,32_ = 14.103, *p* < 0.0001; LHbM: *F*_1,32_ = 32.414, *p* < 0.000001) and a significant interaction between both factors (LHbL: *F*_2,32_ = 3.39, *p* < 0.05; LHbM: *F*_2,32_ = 4.37, *p* < 0.05). The most stricking findings are: in both control groups [(PBS or hM4)-Veh and PBS-CNO], restraint induced c-Fos expression only in the LHbM [(PBS or hM4)-Veh, *p* < 0.05; PBS-CNO, *p* < 0.05], whereas in the hM4-CNO group restraint induced a significant increase in c-Fos expression not only in the LHbM (*p* < 0.001) but also in the LHbL (*p* < 0.001); in the HC condition, c-Fos expression was higher in the LHbM of the hM4-CNO group in comparison to both control groups (*p* < 0.05 for each comparison). These results confirm that stress induces activation predominantly of the medial subdivision of the LHb as already shown in the literature (e.g. Chastrette et al., 1991; Wirtshafter et al., 1994; Brown and Shepard, 2013; Durieux et al., 2020). They also surprisingly show a marked increased activation the LHb of both hM4-CNO groups. To investigate this, we performed fluorescent immunohistochemical staining of the c-Fos protein and performed counting of the number of cells presenting c-Fos+ or/and mCherry+ staining within the LHb. Results are presented in **Figure 3D-F** and show that there are very few cells presenting a co-staining of both c-Fos and mCherry in comparison with c-Fos+ or mCherry+ cells. In fact, in both the hM4-CNO-HC and the hM4-CNO-RS groups, doubly labeled neurons represented only about 1% of c-Fos+ cells (0.95 ± 0.23 % and 1.05 ± 0.29 %, respectively), demonstrating that DREADDed neurons and c-Fos+ neurons were mainly two segregated populations. Such an increased activation of non-DREADDed neurons of the structure could reflect feedback mechanisms from structures downstream to the LHb and influenced by it.

### Functional Network activated by Restraint in (PBS or hM4)-Veh-RS animals

We first investigated if the LHb had a significant position within the stress response control network coming out of the analysis in animals not exposed to CNO, i.e., in (PBS or hM4)-Veh-RS rats. To this purpose, we first calculated the correlation matrix including correlations between all structures, as an estimation of functional connectivity (**Figure 4A**). This highlighted several interesting correlations. For example, we found high connectivity within local networks, such as the mPFC, the AMG and the midbrain monoaminergic regions. This makes perfect sense because those structures are largely described to be crucial in the stress response. We also found high correlations between structures of those different networks, suggesting covariation of activities of cortical, amygdalar and monoaminergic structures upon stress. In other important point looking at the correlation matrix, is that it presents these pools of structures jointly variating called modules. These modules represent complex and robust networks. To test the randomness of these modules, we compared using classical bootstrap (resampled with replacement – 500 repetitions) the (PBS or hM4)-Veh-RS matrix with random matrices created from the bootstrapped (PBS or hM4)-Veh-RS matrices. The comparison of the modularity of both matrices shows a significantly lower modularity coefficient in the random matrix than in the initial data (bootstrap, *p* < 0.05; **Figure 4B**), suggesting that the modularity observed is not random; this allows us to investigate the composition of the different modules and the inter-structural interactions (see later).

**FIGURE 4.**
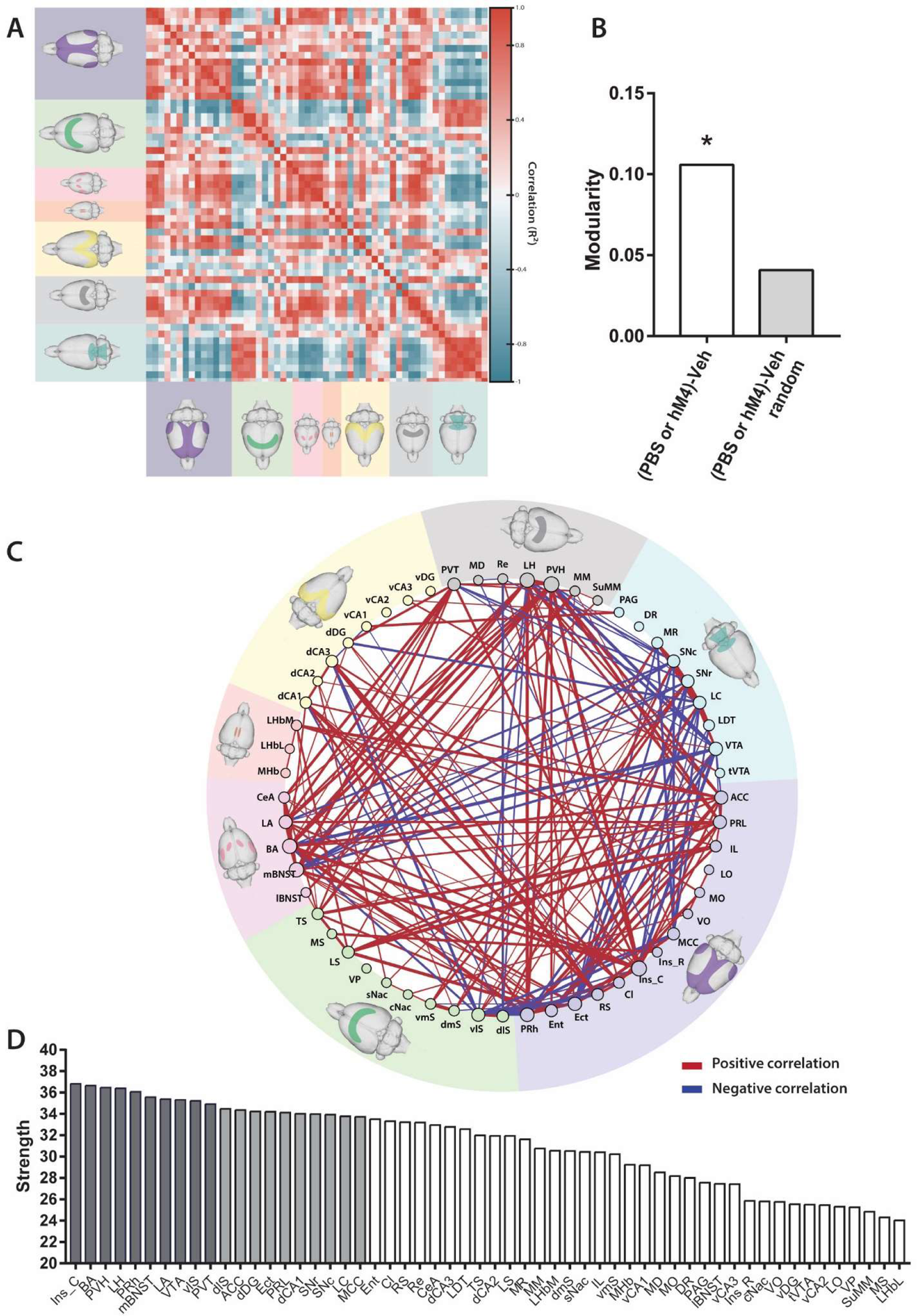
Evaluation of the network engaged by restraint. (**A**) Heatmap representing the cross-correlations (color scale, Pearson correlation R²; red represent positive, and blue negative, correlations) between all investigated structures represented with a color code according to main brain regions (cortical regions, purple; basal ganglia and septum, green; extended amygdala, pink; habenula, red; hippocampus, yellow; thalamus and hypothalamus, grey; brainstem, blue). (**B**) Modularity based on Louvain algorithm, in the (PBS or hM4)-Veh-RS group and its associated random network. <u>Statistics</u>: bootstrap, \**p* < 0.05. (**C**) Graphical representation of the network with nodes (structures) and edges (connecting lines representing significant correlations; Pearson *p* < 0.05). Positive correlations are represented in red and negative ones in blue. (**D**) Strength of each structure in decreasing order. The ten structures (Ins_C to PVT) displaying the highest strength are represented in dark grey and the tens following in light grey.

Significant correlation (Pearson; *p* < 0.05) are represented on a graph theory network (**Figure 4C**). In this network, showing the functional connectivity shared by the structures investigated, we can again notice a large network which is mainly supported by cortical areas (at the exception of the orbitofrontal cortex), the extended amygdala and some thalamic and hypothalamic regions, such as the periventricular nuclei. Furthermore, we identified hubs of connectivity in this network following calculation of the strength of each structure (**Figure 4D**); those key structures include: the Ins_C, the BLA, the PRh, the PVH, the LH, the mBNST, the VTA, the vlS, and the PVT. The analysis indicates that these structures are those including the highest density of shared connections within the network, suggesting that a large part of the flow of information that passes within the network during stress exposure transits through those structures. If we go further in the analysis, the 10 next structures which show a smaller – but still significant – amount of connections, include structures known for their implication in the stress response, such as the anterior and mid cingulate cortices, the dorsal hippocampus (dCA1 and dDG), and monoaminergic regions, *i.e.,* the dopaminergic SNc and noradrenergic LC. According to our main structure of interested, the LHb, whereas its medial subdivision (LHbM) is situated in the medial portion of the graph, making it a hub of medium importance, the LHbL is the last structure that comes out, suggesting it is a very weak actor in the stress response.

To better understand the place of the LHb within the network, we focused our analysis on its correlations with the rest of it (**Figure 5A**). We found several significant correlations (*p* < 0.05 for each association, *R*² > 0.81) with the entire mPFC (ACC; PRL; IL), the LS, and the MHb. As those regions directly project onto the LHb, these results suggest that during stress exposure the LHb process selective information directly coming from those areas. Knowing that the network is composed of non-random modules, we calculated the probability of two structures to be in the same modules across bootstrap repetitions (500 iterations), considering each possible pair of structures (allegiance); further, we focused on the composition of the module that includes the LHbM, as well as the position of this structures within the module.

**FIGURE 5.**
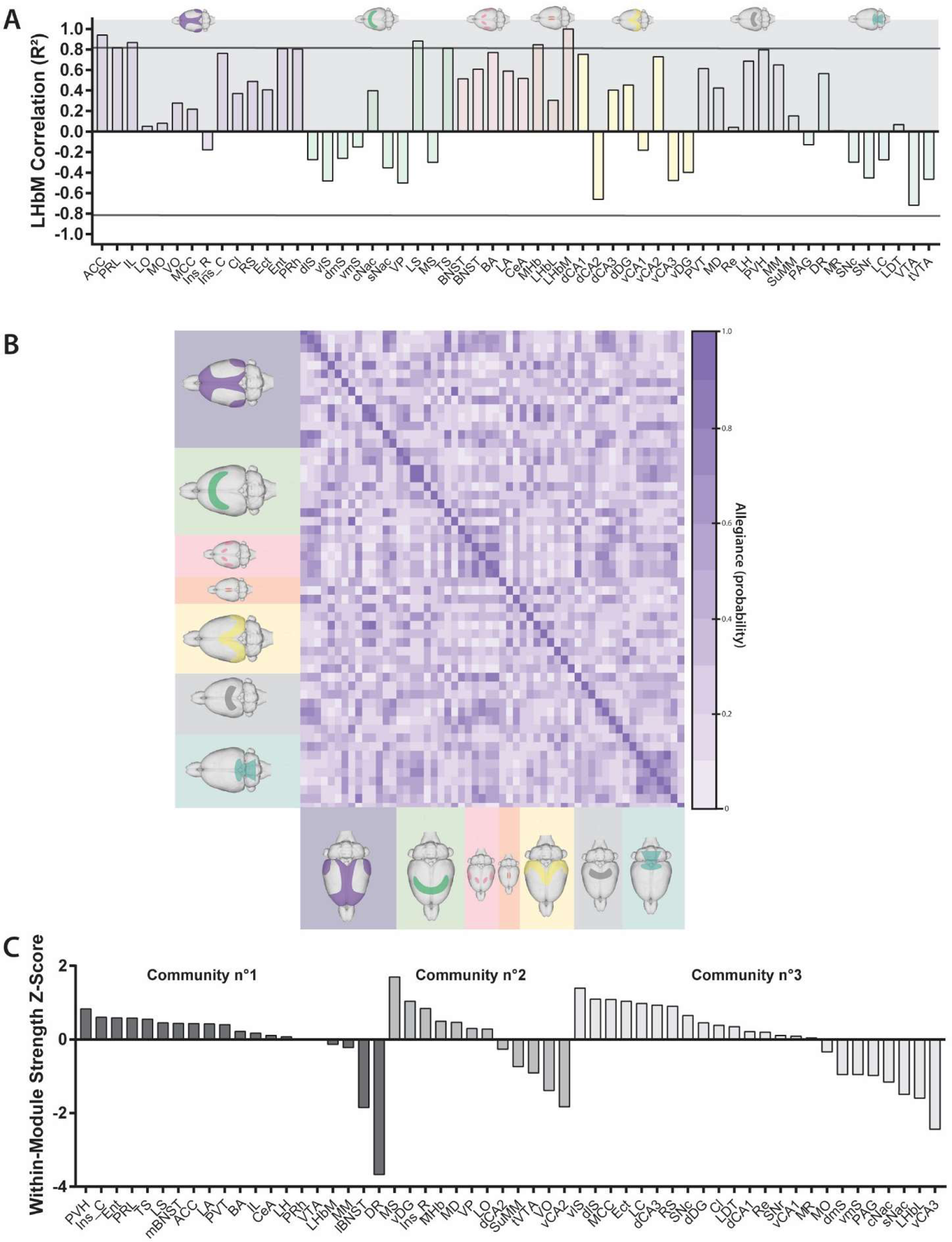
LHbM implication in the stress response may be supported by its functional connections. (**A**) Pearson correlation (*R*²) between the LHbM and structures of the network. Horizontal black lines represent the threshold of significant correlations (*p* < 0.05). The structures are ordered by areas (cortical areas, purple; basal ganglia and septum, green; extended amygdala, pink; habenula, red; hippocampus, yellow; thalamus and hypothalamus, grey; brainstem, blue). Note that the correlation of the LHbM with itself equals 1. (**B**) Heatmap of allegiance of the (PBS or hM4)-Veh-RS group, representing the probability for two given structures to belong to the same community across the bootstrap iterations (the darker, the higher probability). (**C**) Within-modules strength z-score organized by structures according to the community they belong to. The communities have been detected on the allegiance matrix of the (PBS or hM4)-Veh-RS with Louvain algorithm.

The allegiance heatmap displayed three modules, also called “communities” (**Figure 5B**). The community including the LHbM also contained the whole mPFC (ACC, PRL, IL), the Ins_C, the Ent, the PRh, the LS, the TS, both lateral and medial subdivisions of the BNST, the whole AMG, the PVT, the LH, the PVH, the MM, the RD, and the VTA. The community within-module strength Z-score, provides the strength of the connections that a given structure shares with structures of the same community or with structures of other communities (i.e., strong positive scores represent high interaction with structures of the same community, and strong negative scores represent high interaction with structures of other communities, whereas scores around 0 indicate potential bi-directional interactions, intra- and extra-community) (**Figure 5C**). Community n°1, which includes the LHbM, presents an unbalanced pattern of intra/extra community interactions; indeed, only two structures, the lBNST and the RD, seem to connect with structures of different communities, whereas the other structures seem to either show a preference for internal communication or do not show any preference. Communities n°2 and n°3 display a more balanced pattern; around half of the structures promote intra-community interactions, the other half promoting extra-community interactions. Interestingly, the structures present in the same community than the LHbM are crucial for the stress response, i.e., the PVH, the PRL, the mBNST, the ACC, and the LA; moreover, they seem to preferentially display intra-community interaction, suggesting they promote a loop for information processing inside their own community. On the other hand, the lBNST and the RD could be seen as input and/or output structures, as they interact with other communities although we cannot know the directionality of these interactions, whether they receive information from other communities to transmit them to their own, or whether they transmit information from their own community to others. According to the LHbM, its within module strength z-scores – near 0 – suggests it could have a role covering this of the other structures of the community, participating to intra-community interactions and/or to communication with other communities either as an input or an output structure.

### Effects of CNO on functional connectivity

The observation of the correlation matrices of the (PBS or hM4)-Veh-RS and the PBS-CNO-RS groups (**Figure 6A-B**) suggested a decreased functional connectivity in the Ctl-CNO-RS group (**Figure 6B**: colors are globally lighter), although cortical areas (**Figure 6B** top left purple color) display a higher intra-correlation state, suggesting a hypersynchrony of those cortical networks. We then evaluated the strength difference between both (PBS or hM4)-Veh-RS and PBS-CNO-RS networks (**Figure 6C**) and found that some structures showed a significantly higher strength within the (PBS or hM4)-Veh-RS group (classical bootstrap *p* < 0.05 and permutation test *p* < 0.05; ACC, dlS, TS, PVT, LH, PVH, VTA), than within the PBS-CNO-RS group. These results suggest CNO on its own might have altered the brain’s functional network so that it might be difficult to evaluate the net effect of the DREADD-induced LHb inactivation.

**FIGURE 6.**
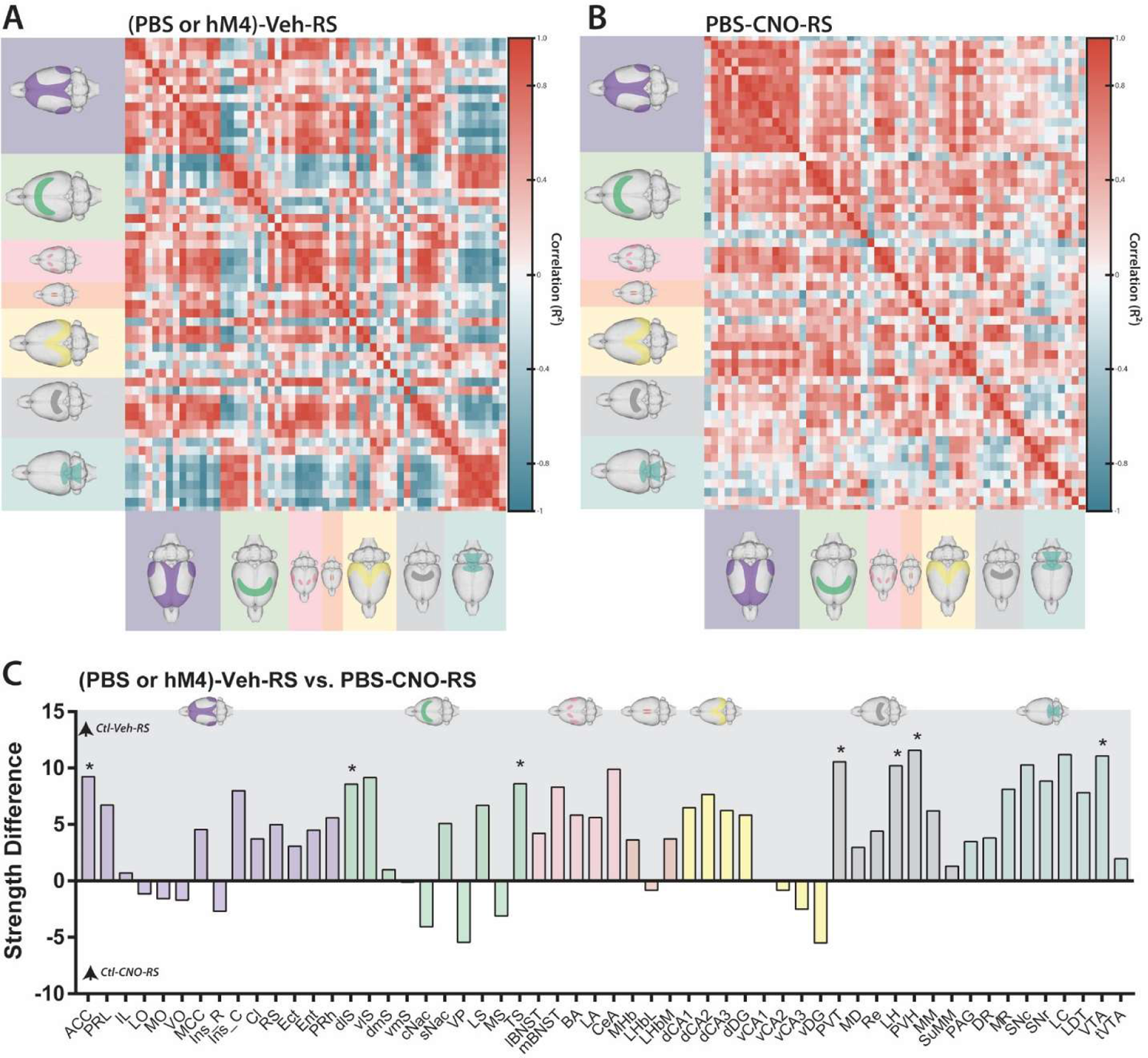
Effects of CNO injection on brain functional network. Heatmap representing the cross-correlations (color scale, Pearson correlation R²; red represent positive, and blue negative, correlations) between all structures organized by main regions (cortical regions, purple; basal ganglia and septum, green; extended amygdala, pink; habenula, red; hippocampus, yellow; thalamus and hypothalamus, grey; brainstem, blue) for (PBS or hM4)-Veh-RS (**A**) and PBS-CNO-RS (**B**). (**C**) Strength difference for each structure between the (PBS or hM4)-Veh-RS group and the PBS-CNO-RS group organized by main areas (positive scores indicate a higher strength in the (PBS or hM4)-Veh-RS group compared to PBS-CNO-RS group; negative scores indicate a higher strength in the PBS-CNO-RS than in (PBS or hM4)-Veh-RS group). <u>Statistics</u>: bootstrap \**p* < 0.05 between groups.

We also investigated the potential effect of CNO based on the stress response control network (represented by the network of the (PBS or hM4)-Veh-RS group), meaning on the strength of the initial communities [communities extracted from the (PBS or hM4)-Veh-RS group]. Accordingly, the correlation matrix of the PBS-CNO-RS group has been sorted according to the structural configuration of the communities of the (PBS or hM4)-Veh-RS group. We were able to visually notice the disruption of (PBS or hM4)-Veh-RS communities in the PBS-CNO-RS group (**Figure 7A** right). In addition, the evaluation of community strength across both networks [(PBS or hM4)-Veh-RS and PBS-CNO-RS] showed that community n°1 and community n°3 were affected by CNO; indeed, the presence of CNO decreased the strength of both communities in comparison with the (PBS or hM4)-Veh group (Mann-Whitney non-parametric test; (PBS or hM4)-Veh-RS *vs.* PBS-CNO-RS for each community; *p* < 0.05; **Figure 7B**). The fact that community n°1, which contains some of the main structures implicated in the stress responses, has a lower strength in the PBS-CNO-RS network, further strengthens the view that this response may have been altered by the presence of CNO.

**FIGURE 7.**
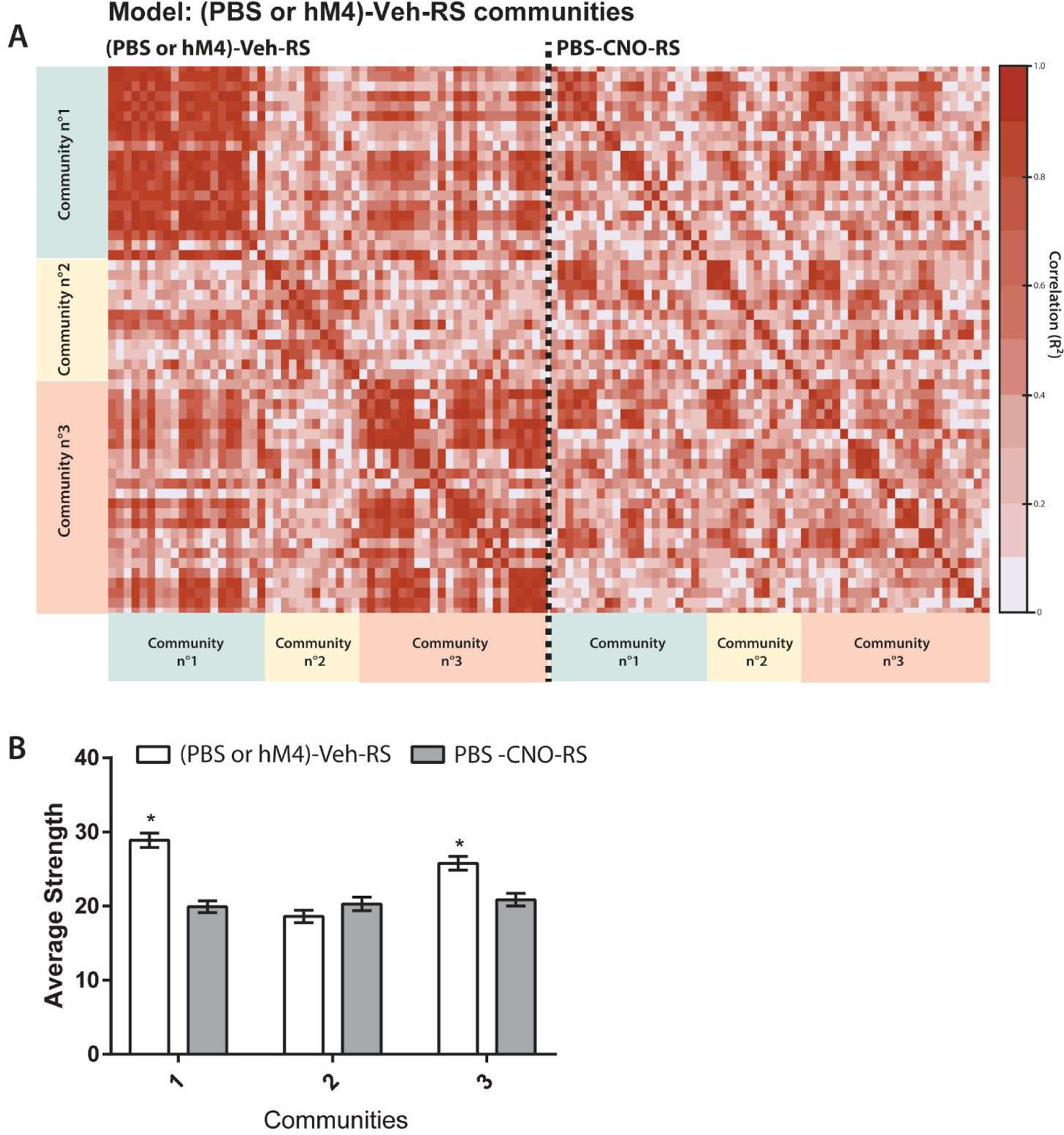
Effects of CNO injection network communities upon restraint. (**A**) Heatmaps representing the cross-correlations (color scale, Pearson correlation R²; the darker, the stronger) between all structures organized according to (PBS or hM4)-Veh-RS communities (communities have been detected using the allegiance of the (PBS or hM4)-Veh-RS group) for both groups [(PBS or hM4)-Veh-RS and PBS-CNO-RS]. (**B**) Average strength of the communities in (PBS or hM4)-Veh-RS and PBS-CNO-RS group. <u>Statistics</u>: Mann-Whitney non-parametric test; \**p* < 0.05 vs. PBS-CNO for the same community.

### Effects of LHb inactivation on functional connectivity

The observation of the correlation matrices of the PBS-CNO-RS and hM4-CNO-RS groups suggests that the heatmaps of both groups are not identical but very similar (**Figure 8A**). We found several significant differences regarding the strength of the structures between the (PBS or hM4)-Veh-RS and PBS-CNO-RS and hM4-CNO-RS, but no difference between the PBS-CNO-RS and the hM4-CNO-RS groups (see Supporting Information **Table S8**), suggesting that the strengths within the network may mainly be the consequence of CNO itself, masking the effects of LHb inactivation. The hM4-CNO-RS group had a quite similar community correlation heatmap than the PBS-CNO group when arranged according to PBS-CNO-RS communities (**Figure 8B**). The allegiance analysis detected three communities in the PBS-CNO-RS group (see Supporting Information, Composition of the communities for each group). The calculation of the average strength of the communities revealed that the strength of community n°1 and n°2 was significantly higher in hM4-CNO-RS group than in PBS-CNO-RS (*p* < 0.05; **Figure 8C**), supporting the idea of an increased community synchrony in animals with LHb inactivation.

**FIGURE 8.**
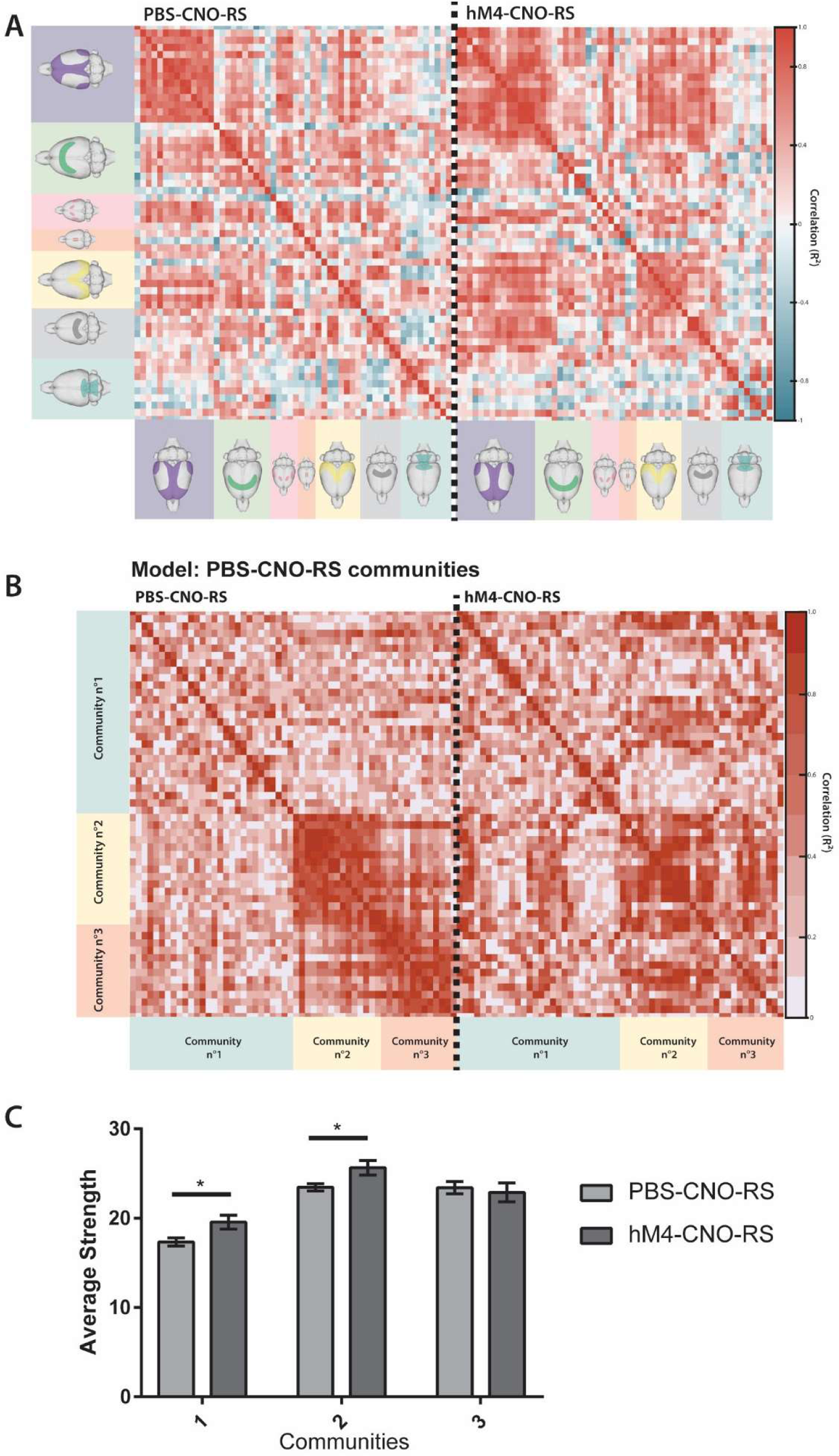
LHb Inactivation and CNO injection alone present some similarities. (**A**) Heatmap representing the cross-correlations (color scale, Pearson correlation *R*²; red represent positive, and blue negative, correlations) between all structures sorted by main regions (cortical areas, purple; basal ganglia and septum areas, green; extended amygdala, pink; habenula, red; hippocampus field, yellow; thalamus and hypothalamus area, grey; brainstem area, blue) for PBS-CNO-RS and hM4-CNO-RS. (**B**) Heatmaps of the correlations (*R*²; the darker the stronger correlation) in PBS-CNO-RS and hM4-CNO-RS groups organized according to PBS-CNO-RS communities (communities extracted on the allegiance matrix of the PBS-CNO-RS group). (**C**) Average strength of the communities in PBS-CNO-RS and hM4-CNO-RS group. <u>Statistics</u>: Mann-Whitney non-parametric test; \**p* < 0.05 indicated between-group difference for the same community.

In addition, we have extracted the composition of the communities of each group in order to compare them (after was checked their “non-randomness” and was calculated their allegiance matrices; see Supporting Information **Figure S6-8**). We calculated the Jaccard coefficient for every possible pair of communities [(PBS or hM4)-Veh-RS vs PBS-CNO-RS, **Figure 9A**; PBS-CNO-RS vs hM4-CNO-RS, **Figure 9B**; details on the composition of each community of each group is given at the end of the Supporting Information]. The composition of the communities of the (PBS or hM4)-Veh-RS group does not show much difference with this of the PBS-CNO group. For example, communities n°1 and n°3 of the (PBS or hM4)-Veh-RS group network are best represented by community n°1 of the PBS-CNO-RS group network (31.4 % and 31.6 % respectively). This suggests that communities n°1 and 3 of the (PBS or hM4)—Veh-RS group have been shuffled in the PBS-CNO-RS group, leading to a reorganization of the balance between the initial communities [i.e. the communities of the (PBS or hM4)-Veh-RS group, based on the original assumption that CNO should not have had any effect and that communities of the PBS-CNO group should have been the same than those of the (PBS or hM4)-Veh-RS group] under the influence of CNO. Also, there are more correspondences when comparing communities of the PBS-CNO-RS and hM4-CNO-RS groups. For example, community n°1 of the PBS-CNO-RS group is highly represented in community n°2 of the hM4-CNO-RS group (47.1 %), suggesting that a large part of the consequences of LHb inactivation on the functional network engaged in the stress response might be due to CNO itself. These results can be used as another indication of the non-negligible consequences of CNO administration.

**FIGURE 9.**
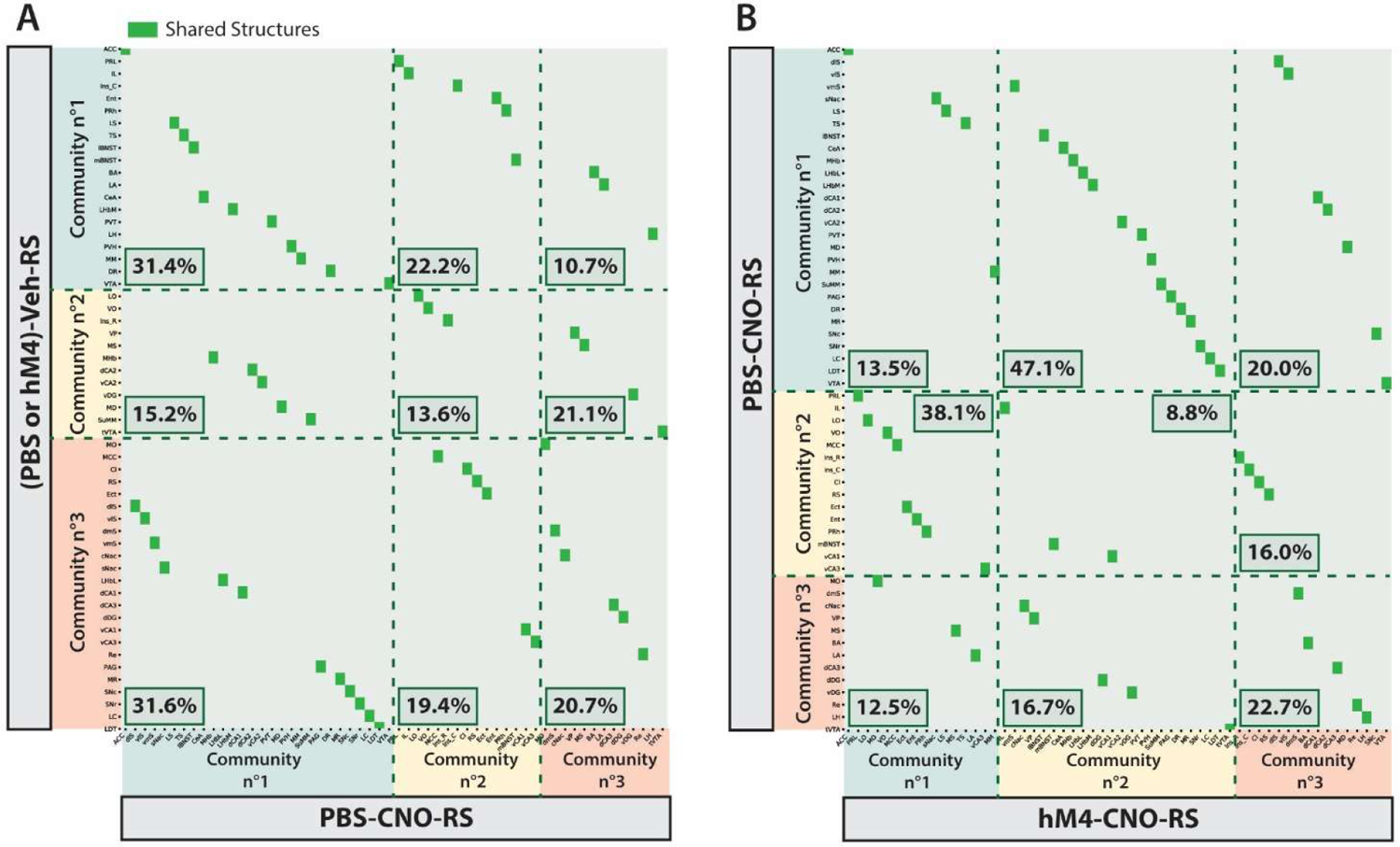
Representation of the structures shared between communities of the different groups. Shared structures (green squares) between (PBS or hM4)-Veh-RS communities and PBS-CNO-RS communities (**A**) and between PBS-CNO-RS communities and hM4-CNO-RS communities (**B**). Jaccard index in percentage is displayed for each comparison, representing the percentage of corresponding between the two communities indicated (rows and columns).

## DISCUSSION

Restraint has extensively been used in the literature as a model of acute stress (Paré and Glavin, 1986). The increased plasmatic CORT concentration observed in all groups seems to validate, if needed, this stressful procedure, as it stimulates the HPA axis. The lack of impact of LHb inactivation on CORT release appears consistent with those already found in our laboratory, showing that LHb inactivation only seems to affect CORT release when there is a cognitive task to perform (Mathis et al., 2018).

Following restraint, we found a generalized marked increase in the density of c-Fos+ cells, indicating the engagement of many different brain structures in the stress response, as expected (Herman et al., 2003; Sousa, 2016). The functional network that seems to support this response possesses a significant modularity (communities of structures which vary together). This modular architecture is a characteristic of small world networks (i.e., a high number of short connections between structures and a high level of local clustering between structures). This type of network is known to be very robust to structural alterations (if one of the nodes is deleted, the information that originally should have passed through this node is eventually rerouted to finally reach its destination without increasing too much the distance it travelled; Sporns et al. 2004; Bassett and Bullmore 2006; Stam and Reijneveld 2007; Reijneveld et al. 2007). Small world architectures are known to be robust networks, promoting rapid information transfer and network synchronization, and providing efficient balance between global integration and local processing (Sporns, 2013). Accordingly, the network engaged upon restraint may be very plastic and robust permitting a very integrated balance of local and global processing leading to highly efficient information transfer at low energy cost.

We found the principal hubs of the stress response network to be the Ins_C, the BA and the PVH, which are main components of the stress response network. The Ins is associated with the modulation of the autonomic response, including somatosensory, nociceptive and visceral processing (Kimura et al., 2010; Menon and Uddin, 2010; Rogers-Carter et al., 2018; Aguilar-Rivera et al., 2020; Ming et al., 2020); in addition, the more caudal part of the Ins (Ins_C), which is the one coming out in our analysis, is involved in respiratory and cardiovascular controls (Bagaev and Aleksandrov, 2006). The engagement of the Ins_C makes perfect sense as the autonomic response facing stress is crucial to allow efficient coping (Selye, 1950; de Kloet et al., 1998). The PVH and the BA are also key structures of the stress response (Prewitt and Herman, 1998; Roozendaal et al., 2009; Sousa, 2016). As said above, the PVH is the entry point of the HPA axis which triggers the physiological response to a stressor through the stimulation of CORT release (Herman et al., 2003; Ulrich-Lai and Herman, 2009). The BLA is essential for processing stressors (Janak and Tye, 2015); it is activated by the anticipation of a stressor (Cullinan et al., 1995) and involved in the consolidation of aversive memories (Roozendaal et al., 2009). Other key structures of the stress response were present in the twenty first hubs that came out from the analysis, including the mBNST, the LA, the mPFC (ACC, PRL), the HPC (dDG, dCA1), and the dopaminergic (SNc and VTA) and noradrenergic (LC) systems (Herman et al., 2003; Sousa, 2016; Godoy et al., 2018).

With regard to the structure of interest of the current study, the LHb, our results first confirmed that its medial part (LHbM) is preferentially activated by stress (Chastrette et al., 1991; Wirtshafter et al., 1994; Brown and Shepard, 2013; Durieux et al., 2020). The LHbM has long been referred to as the “limbic” subdivision of the LHb, as it is directly connected with the mPFC, but also with the hypothalamus and the monoamine systems (Metzger et al., 2021). It therefore makes perfect sense that in the subsequent community analyses of the (PBS or hM4)-Veh-RS group we found strong correlation between the LHbM and the mPFC (ACC, PRL and IL). In addition, we found the LHbM to be significantly correlated to the LS and the MHb. The proximity with the LS is not surprising as identical correlation has been demonstrated using PET + 18-FDG in rats exposed to inescapable foot-shocks (Mirrione et al., 2014). The link with the MHb is more surprising but very interesting. Surprising because the current view is generally that the LHb and the MHb are distinct in terms of anatomical connections and functions; first of all, although the two habenular subregions are indeed very distinct in terms of input and output connections, Kim and Chang (2005) have described a small subset of MHb neurons making en passant synapses on dendrites of LHb neurons before exiting the MHb by the fasciculus retroflexus, suggesting that both regions can share information; these information can indeed be related to the stress response as molecular mechanisms within the MHb have recently been involved in despair-like behavior in mice (Yoo et al., 2021) whereas c-Fos expression was found to be increased in the MHb in rats susceptible to chronic mild stress (Febbraro et al., 2017). These different results imply that both the LHb and MHb could conjointly participate to stress-related responses, such as suggested by our own findings.

Community analyses revealed that, in addition to the mPFC, the LHbM shared strong functional connections with the PVH, the Ins_C, the extended amygdala (BA, LA, CeA, BNST), and the dopamine (VTA) and serotonin (RD) systems. These results support the view that the LHb receives and integrates information from several macrosystems (Geisler and Trimble, 2008), including cortical and amygdalar, related to several aspects of the stress response, and that it could represent a relay of those information towards the monoaminergic centers (VTA and RD), which are engaged in the control of the activity of cortical and subcortical structures to favor coping.

An important result of our study that should be highlighted for future studies is the unexpected effect of CNO on the stress network. Behavioral effects of CNO have generally been observed following administration of doses of CNO higher than the one we used (Manvich et al., 2018), while the vast majority of the studies did not report any adverse behavioral consequences following CNO administration (*e.g.,* Aponte et al., 2011; Ferguson et al., 2013; Augur et al., 2016; Han et al., 2017; Durieux et al., 2020). Our analyses demonstrate an undeniable effect of CNO as correlation matrices of (PBS or hM4)-Veh-RS and PBS-CNO-RS groups hardly differ. Because in our protocol CNO was injected 30 min before restraint, this can suggest the integrity of the stress response network “to be engaged” during restraint had been compromised in rats of the group including LHb inactivation. These effects are probably related to the metabolism of CNO into clozapine (Gomez et al., 2017; Manvich et al., 2018), a psychoactive compound which can influence many neurotransmitter systems through its action on dopamine (D1 and D4), serotonin (including 5HT1A and 5HT2A), noradrenaline (alpha 1 and 2), histamine (H1), and muscarinic (M1, M3 and M4) receptors (Aringhieri et al., 2018). Importantly, in HC condition, CNO did not induce c-Fos expression in all the brain areas known to be activated by clozapine like the mPFC and the NAc (see *e.g.,* Robertson and Fibiger, 1992); this suggests that although CNO was converted into clozapine, its level was probably too low to induce a generalized effect. However, even if CNO altered the level of c-Fos expression in only a few regions, network analyses indicated a lower strength of many structures following its administration, including the ACC, the PVH, and the VTA. We can therefore hypothesize that the detrimental effects of CNO on the organization of the network came from the perturbation of the modulatory function of the HPA axis and of the dopamine system, two very important modulators of the physiological stress response. Interestingly, in rats, at the same concentration we used (1 mg/kg), CNO reduced the acoustic startle reflex (MacLaren et al., 2016) and induced interoceptive effects that partially substitute to that of a low dose of clozapine (Manvich et al., 2018), suggesting it can affect components of the stress response. In trying to differentiate between the effects of CNO itself and the consequences of LHb inactivation, we could only observe that even if some communities were showing some differences between both conditions, the interactions within the communities of the two groups remained quite similar, suggesting the effect of LHb inactivation had been partly masked by the effects of CNO.

## CONCLUSIONS

We have seen that the LHb, and more specifically its medial subdivision, the LHbM, was engaged, during acute stress exposure, among a broad network of key regions of the stress response. Given its anatomical position, the role of the LHbM could be to integrate multiple stress-related information stemming in cortical, thalamic and hypothalamic regions to further communicate it downstream to monoaminergic systems in order to probably initiate coping strategies. As shown by others, the LHb – or its equivalent in zebrafishes – is engaged in behavioral adaptation under stressful situations (Ootsuka and Mohammed, 2015; Berger et al., 2018; Andalman et al., 2019; Coffey et al., 2020). Better understanding the engagement of the LHb when individuals are exposed to stressful situation appears like a relevant question as alteration of LHb function or morphology has been evidenced in stress-related psychiatric pathologies such as depression (Browne et al., 2018; Gold and Kadriu, 2019; Hu et al., 2020) and schizophrenia (Sandyk, 1992; Shepard et al., 2006; Zhang et al., 2017; but see Schafer et al., 2018).

## List of Abbreviations

AAV8: Adeno-Associated Virus serotype 8

ACC: Anterior Cingulate Cortex

ACN: acetonitrile

AMG: Amygdala

BA: Basal nucleus of the Amygdala

BLA: Basal Lateral nucleus of the Amygdala

BNST: Bed Nucleus of the *Stria Terminalis*

CamKII: CA^2+^/Calmodulin-dependent protein Kinase II

CeA: Central nucleus of the Amygdala

Cl: Claustrum

cNAc: core of the Nucleus Accumbens

CNO: Clozapine-N-Oxide

CORT: Corticosterone

dCA1: dorsal *Cornus Ammonis 1*

dCA2: dorsal *Cornus Ammonis 2*

dCA3: dorsal *Cornus Ammonis 3*

dDG: dorsal Dentate Gyrus

dlS: dorso-lateral Striatum

dmS: dorso-median Striatum

DMSO: Dimethyl sulfoxide

DR: Raphe Dorsal

DREADD: Designer Receptor Exclusively Activated by Designer Drugs

Ect: Ectorhinal cortex

Ent: Entorhinal cortex

hM4(Gi): modified human Muscarinic 4 (coupled with inhibitory G protein)

HPA: hypothalamus-pituitary-adrenal

HPC: Hippocampus

HPLC: High performance liquid chromatography

IL: Infralimbic cortex

Ins_C: Caudal part of the Insular cortex

Ins_R: Rostral part of the Insular cortex

LA: Lateral nucleus of the Amygdala

lBNST: Bed Nucleus of the *Stria Terminalis* (lateral part)

LC: Locus Coeruleus

LDT: Latero-dorsal Tegmental nucleus

LH: Lateral Hypothalamus

LHb: Lateral Habenula

LHbL: Lateral part of the Lateral Habenula

LHbM: Medial part of the Lateral Habenula

LO: Lateral Orbitofrontal cortex

LS: Lateral Septum

mBNST: Bed Nucleus of the *Stria Terminalis* (medial part)

MCC: Mid Cingulate Cortex

MD: Medio-Dorsal nucleus of the thalamus

MHb: Medial Habenula

MM: Mammillary nucleus of hypothalamus

MO: Medial Orbitofrontal cortex

mPFC: medial prefrontal cortex

MR: Raphe Median

MS: Medial Septum

MSpect: Mass Spectrometer

PAG: Periaqueductal Gray

PBS: Phosphate Buffered Saline

PFC: Prefrontal Cortex

PRh: Perirhinal cortex

PRL: Prelimbic cortex

PVH: Paraventricular nucleus of the Hypothalamus

PVT: Paraventricular nucleus of the Thalamus

Re: Reuniens nucleus

RS: Retrosplenial Cortex

sNAc: shell of the Nucleus Accumbens

SNc: *Substantia Nigra pars compacta*

SNr: *Substantia Nigra pars reticula*

SRM: Selected Reaction Monitoring

SuMM: Supramammillary nucleus of the hypothalamus

TS: Triangular Septal nucleus

tVTA: tail of the Ventral Tegmental Area **v**

CA1: ventral *Cornus Ammonis* 1

vCA2: ventral *Cornus Ammonis* 2

vCA3: ventral *Cornus Ammonis* 3

vDG: ventral Dentate Gyrus

vlS: ventro-lateral Striatum

vmS: ventro-median Striatum

VO: Ventral Orbitofrontal cortex

VP: Ventral Pallidum

VTA: Ventral Tegmental Area

## ACKNOWLEDGEMENTS

This research was supported by the CNRS, Université de Strasbourg, and a ERA-NET NEURON grant to LL (Neuromarket). The authors wish to thank Dr. Michel Barrot for the gift of CNO.

## CONFLICT OF INTEREST

The authors declare no conflict of interest.

## ETHICAL STATEMENT

The authors declare that animals were handled in accordance with ethical requirements enacted by the French authorities and the Strasbourg university who approved the present project (APAFIS #7114).

## AUTHOR CONTRIBUTIONS

LL and MM designed the study. LD, LL, MM performed the experiments and analysed the data. AB helped with the restraint procedure. KH, CB, CH helped with the c-fos counting. DB helped with graph theory analyses. VA and YG performed the LC-MS/MS measurements. LL, MM, LD wrote the manuscript.

## DATA AVAILABILITY STATEMENT

Data supporting the findings of this study are available upon reasonable request from the corresponding authors.

## SUPPORTING INFORMATION

Additional supporting information may be found online in the Supporting Information section at the end of this article.

## Notes

### Competing Interest Statement

The authors have declared no competing interest.

## REFERENCES

1. Aguilar-Rivera, M., Kim, S., Coleman, T.P., Maldonado, P.E., Torrealba, F. (2020). Interoceptive insular cortex participates in sensory processing of gastrointestinal malaise and associated behaviors. Sci Rep, 10, 21642.

2. Aizawa, H., Yanagihara, S., Kobayashi, M., Niisato, K., Takekawa, T., Harukuni, R., McHugh, T.J., Fukai, T., Isomura, Y., Okamoto, H. (2013). The synchronous activity of lateral habenular neurons is essential for regulating hippocampal theta oscillation. J Neurosci, 33, 8909–21.

3. Ali, M., Cholvin, T., Muller, M.A., Cosquer, B., Kelche, C., Cassel, J.-C., Pereira de Vasconcelos; A. (2017). Environmental enrichment enhances systems-level consolidation of a spatial memory after lesions of the ventral midline thalamus. Neurobiol Learn Mem, 141, 108–123.

4. Andalman, A.S., Burns, V.M., Lovett-Barron, M., Broxton, M., Poole, B., Yang, S.J., Grosenick, L., Lerner, T.N., Chen, R., Benster, T., Mourrain, P., Levoy, M., Rajan, K., Deisseroth, K. (2019). Neuronal Dynamics Regulating Brain and Behavioral State Transitions. Cell, 177, 970–985.

5. Aponte, Y., Atasoy, D., Sternson, S.M. (2011). AGRP neurons are sufficient to orchestrate feeding behavior rapidly and without training. Nat Neurosci, 14, 351–355.

6. Aringhieri, S., Carli, M., Kolachalam, S., Verdesca, V., Cini, E., Rossi, M., McCormick, P.J., Corsini, G.U., Maggio, R., Scarselli, M. (2018). Molecular targets of atypical antipsychotics: From mechanism of action to clinical differences. Pharmacol Ther, 192, 20–41.

7. Augur, I.F., Wyckoff, A.R., Aston-Jones, G., Kalivas, P.W., Peters, J. (2016). Chemogenetic Activation of an Extinction Neural Circuit Reduces Cue-Induced Reinstatement of Cocaine Seeking. J Neurosci, 36, 10174–10180.

8. Bagaev, V., Aleksandrov, V. (2006). Visceral-related area in the rat insular cortex. Auton Neurosci Basic Clin, 125, 16–21.

9. Baker, P.M., Rao, Y., Rivera, Z.M.G., Garcia, E.M., Mizumori, S.J.Y. (2019). Selective Functional Interaction Between the Lateral Habenula and Hippocampus During Different Tests of Response Flexibility. Front Mol Neurosci, 12, 245.

10. Bankhead, P., Loughrey, M.B., Fernández, J.A., Dombrowski, Y., McArt, D.G., Dunne, P.D., McQuaid, S., Gray, R.T., Murray, L.J., Coleman, H.G., James, J.A., Salto-Tellez, M., Hamilton, P.W. (2017). QuPath: Open source software for digital pathology image analysis. Sci Rep, 7, 16878.

11. Barrett, D.W., Gonzalez-Lima, F. (2018). Prefrontal-limbic Functional Connectivity during Acquisition and Extinction of Conditioned Fear. Neuroscience, 376, 162–171.

12. Bassett, D.S., Bullmore, E. (2006). Small-world brain networks. Neurosci Rev J Bringing Neurobiol Neurol Psychiatry, 12, 512–523.

13. Berger, A.L., Henricks, A.M., Lugo, J.M., Wright, H.R., Warrick, C.R., Sticht, M.A., Morena, M., Bonilla, I., Laredo, S.A., Craft, R.M., Parsons, L.H., Grandes, P.R., Hillard, C.J., Hill, M.N., McLaughlin, R.J. (2018). The Lateral Habenula Directs Coping Styles Under Conditions of Stress via Recruitment of the Endocannabinoid System. Biol Psychiatry, 84, 611–623.

14. Brown, P.L., Shepard, P.D. (2013). Lesions of the fasciculus retroflexus alter footshock-induced cFos expression in the mesopontine rostromedial tegmental area of rats. PloS One, 8, e60678.

15. Browne, C.A., Hammack, R., Lucki, I. (2018). Dysregulation of the Lateral Habenula in Major Depressive Disorder. Front Synaptic Neurosci, 7, 10:46.

16. Bullmore, E., Sporns, O. (2009). Complex brain networks: graph theoretical analysis of structural and functional systems. Nat Rev Neurosci, 10, 186–198.

17. Cenci, M.A., Kalén, P., Mandel, R.J., Björklund, A. (1992). Regional differences in the regulation of dopamine and noradrenaline release in medial frontal cortex, nucleus accumbens and caudate-putamen: a microdialysis study in the rat. Brain Res, 581, 217–28.

18. Chastrette, N., Pfaff, D.W., Gibbs, R.B. (1991). Effects of daytime and nighttime stress on Fos-like immunoreactivity in the paraventricular nucleus of the hypothalamus, the habenula, and the posterior paraventricular nucleus of the thalamus. Brain Res, 563, 339–344.

19. Christoph, G.R., Leonzio, R.J., Wilcox, K.S. (1986). Stimulation of the lateral habenula inhibits dopamine-containing neurons in the substantia nigra and ventral tegmental area of the rat. J Neurosci, 6, 613–9.

20. Coffey, K.R., Marx, R.G., Vo, E.K., Nair, S.G., Neumaier, J.F. (2020). Chemogenetic inhibition of lateral habenula projections to the dorsal raphe nucleus reduces passive coping and perseverative reward seeking in rats. Neuropsychopharmacol Off Publ Am Coll Neuropsychopharmacol, 45, 1115–1124.

21. Cullinan, W.E., Herman, J.P., Battaglia, D.F., Akil, H., Watson, S.J. (1995). Pattern and time course of immediate early gene expression in rat brain following acute stress. Neuroscience, 64, 477–505.

22. de Kloet, E.R., Vreugdenhil, E., Oitzl, M.S., Joëls, M. (1998). Brain Corticosteroid Receptor Balance in Health and Disease. Endocr Rev, 19, 269–301.

23. Derman, R.C., Bass, C.E., Ferrario, C.R. (2020). Effects of hM4Di activation in CamKII basolateral amygdala neurons and CNO treatment on sensory-specific vs. general PIT: refining PIT circuits and considerations for using CNO. Psychopharmacology (Berl), 237, 1249–1266.

24. Dolzani, S.D., Baratta, M.V., Amat, J., Agster, K.L., Saddoris, M.P., Watkins, L.R., Maier, S.F. (2016). Activation of a Habenulo-Raphe Circuit Is Critical for the Behavioral and Neurochemical Consequences of Uncontrollable Stress in the Male Rat. eNeuro, 3.

25. Dong, H.W., Swanson, L.W. (2004). Organization of axonal projections from the anterolateral area of the bed nuclei of the stria terminalis. J Comp Neurol, 468, 277–98.

26. Dong, H.W., Swanson, L.W. (2006). Projections from bed nuclei of the stria terminalis, dorsomedial nucleus: implications for cerebral hemisphere integration of neuroendocrine, autonomic, and drinking responses. J Comp Neurol, 494, 75–107.

27. Durieux, L., Mathis, V., Herbeaux, K., Muller, M.-A., Barbelivien, A., Mathis, C., Schlichter, R., Hugel, S., Majchrzak, M., Lecourtier, L. (2020). Involvement of the lateral habenula in fear memory. Brain Struct Funct, 225, 2029–2044.

28. Febbraro, F., Svenningsen, K., Tran, T.P., Wiborg, O. (2017). Neuronal substrates underlying stress resilience and susceptibility in rats. PloS One, 12, e0179434.

29. Ferguson, S.M., Phillips, P.E.M., Roth, B.L., Wess, J., Neumaier, J.F. (2013). Direct-Pathway Striatal Neurons Regulate the Retention of Decision-Making Strategies. J Neurosci, 33, 11668– 11676.

30. Geisler, S., Trimble, M. (2008). The Lateral Habenula: No Longer Neglected. CNS Spectr, 13, 484– 489.

31. Godoy, L.D., Rossignoli, M.T., Delfino-Pereira, P., Garcia-Cairasco, N., de Lima Umeoka, E.H. (2018). A Comprehensive Overview on Stress Neurobiology: Basic Concepts and Clinical Implications. Front Behav Neurosci, 12, 127.

32. Gold, P.W., Kadriu, B. (2019). A Major Role for the Lateral Habenula in Depressive Illness: Physiologic and Molecular Mechanisms. Front Psychiatry, 10, 320.

33. Gomez, J.L., Bonaventura, J., Lesniak, W., Mathews, W.B., Sysa-Shah, P., Rodriguez, L.A., Ellis, R.J., Richie, C.T., Harvey, B.K., Dannals, R.F., Pomper, M.G., Bonci, A., Michaelides, M. (2017). Chemogenetics revealed: DREADD occupancy and activation via converted clozapine. Science, 357, 503–507.

34. González-Pardo, H., Conejo, N.M., Lana, G., Arias, J.L. (2012). Different brain networks underlying the acquisition and expression of contextual fear conditioning: a metabolic mapping study. Neuroscience, 202, 234–242.

35. Goutagny, R., Loureiro, M., Jackson, J., Chaumont, J., Williams, S., Isope, P., Kelche, C., Cassel, J.C., Lecourtier, L. (2013). Interactions between the lateral habenula and the hippocampus: implication for spatial memory processes. Neuropsychopharmacology, 38, 2418–26.

36. Han, S., Yang, S.H., Kim, J.Y., Mo, S., Yang, E., Song, K.M., Ham, B.-J., Mechawar, N., Turecki, G., Lee, H.W., Kim, H. (2017). Down-regulation of cholinergic signaling in the habenula induces anhedonia-like behavior. Sci Rep, 7, 900.

37. Henckens, M.J.A.G., van der Marel, K., van der Toorn, A., Pillai, A.G., Fernández, G., Dijkhuizen, R.M., Joëls, M. (2015). Stress-induced alterations in large-scale functional networks of the rodent brain. NeuroImage, 105, 312–322.

38. Herman, J.P., Figueiredo, H., Mueller, N.K., Ulrich-Lai, Y., Ostrander, M.M., Choi, D.C., Cullinan, W.E. (2003). Central mechanisms of stress integration: hierarchical circuitry controlling hypothalamo–pituitary–adrenocortical responsiveness. Front Neuroendocrinol, 24, 151–180.

39. Hikosaka, O. (2010). The habenula: from stress evasion to value-based decision-making. Nat Rev Neurosci, 11, 503–513.

40. Hu, H., Cui, Y., Yang, Y. (2020). Circuits and functions of the lateral habenula in health and in disease. Nat Rev Neurosci, 21, 277–295.

41. Hudson, A.E. (2018). Genetic Reporters of Neuronal Activity: c-Fos and G-CaMP6. Methods Enzymol, 603, 197–220.

42. Jacinto, L.R., Mata, R., Novais, A., Marques, F., Sousa, N. (2017). The habenula as a critical node in chronic stress-related anxiety. Exp Neurol, 289, 46–54.

43. Janak, P.H., Tye, K.M. (2015). From circuits to behaviour in the amygdala. Nature, 517, 284–292.

44. Ji, H., Shepard, P.D. (2007). Lateral habenula stimulation inhibits rat midbrain dopamine neurons through a GABA(A) receptor-mediated mechanism. J Neurosci, 27, 6923–30.

45. Joëls, M., Baram, T.Z. (2009). The neuro-symphony of stress. Nat Rev Neurosci, 10, 459–466.

46. Kalén, P., Lindvall, O., Björklund, A. (1989a). Electrical stimulation of the lateral habenula increases hippocampal noradrenaline release as monitored by in vivo microdialysis. Exp Brain Res, 76, 239–45.

47. Kalén, P., Strecker, R.E., Rosengren, E., Björklund, A. (1989b). Regulation of striatal serotonin release by the lateral habenula-dorsal raphe pathway in the rat as demonstrated by in vivo microdialysis: role of excitatory amino acids and GABA. Brain Res, 492, 187–202.

48. Kim, U., Chang, S.-Y. (2005). Dendritic morphology, local circuitry, and intrinsic electrophysiology of neurons in the rat medial and lateral habenular nuclei of the epithalamus. J Comp Neurol, 483, 236–250.

49. Kim, U., Lee, T. (2012). Topography of descending projections from anterior insular and medial prefrontal regions to the lateral habenula of the epithalamus in the rat. Eur J Neurosci, 35, 1253–69.

50. Kimura, A., Imbe, H., Donishi, T. (2010). Efferent connections of an auditory area in the caudal insular cortex of the rat: anatomical nodes for cortical streams of auditory processing and cross-modal sensory interactions. Neuroscience, 166, 1140–1157.

51. Kovács, K.J. (1998). Invited review c-Fos as a transcription factor: a stressful (re)view from a functional map. Neurochem Int, 33, 287–297.

52. Lammel, S., Lim, B.K., Ran, C., Huang, K.W., Betley, M.J., Tye, K.M., Deisseroth, K., Malenka, R.C. (2012). Input-specific control of reward and aversion in the ventral tegmental area. Nature, 491, 212–217.

53. Lecca, S., Trusel, M., Mameli, M. (2017). Footshock-induced plasticity of GABAB signalling in the lateral habenula requires dopamine and glucocorticoid receptors. Synapse, 71.

54. Lecourtier, L., Defrancesco, A., Moghaddam, B. (2008). Differential tonic influence of lateral habenula on prefrontal cortex and nucleus accumbens dopamine release. Eur J Neurosci, 27, 1755–62.

55. MacLaren, D.A.A., Browne, R.W., Shaw, J.K., Krishnan Radhakrishnan, S., Khare, P., España, R.A., Clark, S.D. (2016). Clozapine N-Oxide Administration Produces Behavioral Effects in Long– Evans Rats: Implications for Designing DREADD Experiments. eNeuro, 3.

56. Magalhães, R., Barrière, D.A., Novais, A., Marques, F., Marques, P., Cerqueira, J., Sousa, J.C., Cachia, A., Boumezbeur, F., Bottlaender, M., Jay, T.M., Mériaux, S., Sousa, N. (2018). The dynamics of stress: a longitudinal MRI study of rat brain structure and connectome. Mol Psychiatry, 23, 1998–2006.

57. Manvich, D.F., Webster, K.A., Foster, S.L., Farrell, M.S., Ritchie, J.C., Porter, J.H., Weinshenker, D. (2018). The DREADD agonist clozapine N-oxide (CNO) is reverse-metabolized to clozapine and produces clozapine-like interoceptive stimulus effects in rats and mice. Sci Rep, 8.

58. Mathis, V., Cosquer, B., Barbelivien, A., Herbeaux, K., Bothorel, B., Sage-Ciocca, D., Poirel, V.-J., Mathis, C., Lecourtier, L. (2018). The lateral habenula interacts with the hypothalamo-pituitary adrenal axis response upon stressful cognitive demand in rats. Behav Brain Res, 341, 63–70.

59. McEwen, B.S., Bowles, N.P., Gray, J.D., Hill, M.N., Hunter, R.G., Karatsoreos, I.N., Nasca, C. (2015). Mechanisms of stress in the brain. Nat Neurosci, 18, 1353–1363.

60. Menon, V., Uddin, L.Q. (2010). Saliency, switching, attention and control: a network model of insula function. Brain Struct Funct, 214, 655–667.

61. Metzger, M., Souza, R., Lima, L.B., Bueno, D., Gonçalves, L., Sego, C., Donato, J., Shammah-Lagnado, S.J. (2021). Habenular connections with the dopaminergic and serotonergic system and their role in stress-related psychiatric disorders. Eur J Neurosci, 53, 65–88.

62. Ming, Q., Ma, H., Li, J., Yang, F., Li, J., Liang, J., Li, D., Lin, W. (2020). Changes in autonomic nervous function and influencing factors in a rat insular cortex electrical kindling model. Neurosci Lett, 721, 134782.

63. Mirrione, M.M., Schulz, D., Lapidus, K.A.B., Zhang, S., Goodman, W., Henn, F.A. (2014). Increased metabolic activity in the septum and habenula during stress is linked to subsequent expression of learned helplessness behavior. Front Hum Neurosci, 8.

64. Murphy, C.A., DiCamillo, A.M., Haun, F., Murray, M. (1996). Lesion of the habenular efferent pathway produces anxiety and locomotor hyperactivity in rats: a comparison of the effects of neonatal and adult lesions. Behav Brain Res, 81, 43–52.

65. Ootsuka, Y., Mohammed, M. (2015). Activation of the habenula complex evokes autonomic physiological responses similar to those associated with emotional stress. Physiol Rep, 3, e12297.

66. Ootsuka, Y., Mohammed, M., Blessing, W.W. (2017). Lateral habenula regulation of emotional hyperthermia: mediation via the medullary raphé. Sci Rep, 7, 4102.

67. Paré, W.P., Glavin, G.B. (1986). Restraint stress in biomedical research: a review. Neurosci Biobehav Rev, 10, 339–370.

68. Prewitt, C.M.F., Herman, J.P. (1998). Anatomical interactions between the central amygdaloid nucleus and the hypothalamic paraventricular nucleus of the rat: a dual tract-tracing analysis. J Chem Neuroanat, 15, 173–186.

69. Reijneveld, J.C., Ponten, S.C., Berendse, H.W., Stam, C.J. (2007). The application of graph theoretical analysis to complex networks in the brain. Clin Neurophysiol, 118, 2317–2331.

70. Reisine, T.D., Soubrié, P., Artaud, F., Glowinski, J. (1982). Involvement of lateral habenula-dorsal raphe neurons in the differential regulation of striatal and nigral serotonergic transmission cats. J Neurosci, 2, 1062–71.

71. Robertson, G.S., Fibiger, H.C. (1992). Neuroleptics increase C-FOS expression in the forebrain: Contrasting effects of haloperidol and clozapine. Neuroscience, 46, 315–328.

72. Rogers-Carter, M.M., Varela, J.A., Gribbons, K.B., Pierce, A.F., McGoey, M.T., Ritchey, M., Christianson, J.P. (2018). Insular cortex mediates approach and avoidance responses to social affective stimuli. Nat Neurosci, 21, 404–414.

73. Roozendaal, B., McEwen, B.S., Chattarji, S. (2009). Stress, memory and the amygdala. Nat Rev Neurosci, 10, 423–433.

74. Sandyk, R. (1992). Pineal and habenula calcification in schizophrenia. Int J Neurosci, 67, 19–30.

75. Selye, H. (1950). Stress and the General Adaptation Syndrome. Br Med J, 1, 1383–1392.

76. Shabel, S.J., Proulx, C.D., Trias, A., Murphy, R.T., Malinow, R. (2012). Input to the Lateral Habenula from the Basal Ganglia Is Excitatory, Aversive, and Suppressed by Serotonin. Neuron, 74, 475–481.

77. Schafer, M., Kim J.W., Joseph, J., Xu, J., Frangou S., Doucet G.E. (2018) Imaging Habenula Volume in Schizophrenia and Bipolar Disorder. Front Psychiatry, 9, 456.

78. Shepard, P.D., Holcomb, H.H., Gold, J.M. (2006). Schizophrenia in translation: the presence of absence: habenular regulation of dopamine neurons and the encoding of negative outcomes. Schizophr Bull, 32, 417–21.

79. Sousa, N. (2016). The dynamics of the stress neuromatrix. Mol Psychiatry, 21, 302–312.

80. Sporns, O. (2013). Network attributes for segregation and integration in the human brain. Curr Opin Neurobiol, 23, 162–171.

81. Sporns, O., Chialvo, D.R., Kaiser, M., Hilgetag, C.C. (2004). Organization, development and function of complex brain networks. Trends Cogn Sci, 8, 418–425.

82. Stam, C.J., Reijneveld, J.C. (2007). Graph theoretical analysis of complex networks in the brain. Nonlinear Biomed Phys, 1, 3.

83. Stamatakis, A.M., Stuber, G.D. (2012). Activation of lateral habenula inputs to the ventral midbrain promotes behavioral avoidance. Nat Neurosci, 15, 1105–1107.

84. Ulrich-Lai, Y.M., Herman, J.P. (2009). Neural regulation of endocrine and autonomic stress responses. Nat Rev Neurosci, 10, 397–409.

85. Vetere, G., Kenney, J.W., Tran, L.M., Xia, F., Steadman, P.E., Parkinson, J., Josselyn, S.A., Frankland, P.W. (2017). Chemogenetic Interrogation of a Brain-wide Fear Memory Network in Mice. Neuron, 94, 363–374.e4.

86. Wheeler, A.L., Teixeira, C.M., Wang, A.H., Xiong, X., Kovacevic, N., Lerch, J.P., McIntosh, A.R., Parkinson, J., Frankland, P.W. (2013). Identification of a functional connectome for long-term fear memory in mice. PLoS Comput Biol, 9, e1002853.

87. Wirtshafter, D., Asin, K.E., Pitzer, M.R. (1994). Dopamine agonists and stress produce different patterns of Fos-like immunoreactivity in the lateral habenula. Brain Res, 633, 21–26.

88. Yoo, H., Yang, S.H., Kim, J.Y., Yang, E., Park, H.S., Lee, S.J., Rhyu, I.J., Turecki, G., Lee, H.W., Kim, H. (2021). Down-regulation of habenular calcium-dependent secretion activator 2 induces despair-like behavior. Sci Rep, 11, 3700.

89. Zhang, J., Tan, L., Ren, Y., Liang, J., Lin, R., Feng, Q., Zhou, J., Hu, F., Ren, J., Wei, C., Yu, T., Zhuang, Y., Bettler, B., Wang, F., Luo, M. (2016). Presynaptic Excitation via GABAB Receptors in Habenula Cholinergic Neurons Regulates Fear Memory Expression. Cell, 166, 716–728.

90. Zhang, L., Wang, H., Luan, S., Yang, S., Wang, Z., Wang, J., Zhao H. (2017). Altered Volume and Functional Connectivity of the Habenula in Schizophrenia. Front Hum Neurosci, 11, 636.

91. Zhou W, Jin Y, Meng Q, Zhu X, Bai T, Tian Y, Mao Y, Wang L, Xie W, Zhong H, Zhang N, Luo MH, Tao W, Wang H, Li J, Li J, Qiu BS, Zhou JN, Li X, Xu H, Wang K, Zhang X, Liu Y, Richter-Levin G, Xu L, Zhang Z. (2019). A neural circuit for comorbid depressive symptoms in chronic pain. Nat Neurosci, 22, 1649–1658.

