## Supplementary data for "Functional brain-wide network mapping during acute stress exposure in rats: Interaction between the lateral habenula and cortical, amygdalar, hypothalamic and monoaminergic regions"

#### Materials and Methods

**Table S1:** Treatments received by rats throughout the whole experimental procedures, including the previously published study (Durieux et al., 2020).

According to the present study, the color code is as follows: (PBS or hM4)-Veh-HC (light blue), PBS-CNO-HC (light green), hM4-CNO-HC (light orange), (PBS or hM4)-Veh-RS (dark blue), PBS-CNO-RS (dark green), hM4-CNO-RS (dark orange).

| | | Treatments received (previous study) | | | Treatment received (present study) |
|---|---|---|---|---|---|
| Rats # | Surgery | Plus maze | Fear conditioning | Locomotor activity | During restraint (RS) or in home cage (HC) |
| 112 | PBS | Veh | Veh | Veh | Veh (HC) |
| 117 | PBS | Veh | Veh | Veh | Veh (HC) |
| 136 | PBS | Veh | Veh | Veh | Veh (HC) |
| 100 | hM4-mCherry | Veh | Veh | Veh | Veh (HC) |
| 104 | hM4-mCherry | Veh | Veh | Veh | Veh (HC) |
| 106 | hM4-mCherry | Veh | Veh | Veh | Veh (HC) |
| 114 | PBS | CNO | CNO | CNO | CNO (HC) |
| 119 | PBS | CNO | CNO | CNO | CNO (HC) |
| 121 | PBS | CNO | CNO | CNO | CNO (HC) |
| 122 | PBS | CNO | CNO | CNO | CNO (HC) |
| 125 | PBS | CNO | CNO | CNO | CNO (HC) |
| 130 | PBS | CNO | CNO | CNO | CNO (HC) |
| 133 | PBS | CNO | CNO | CNO | CNO (HC) |
| 134 | PBS | CNO | CNO | CNO | CNO (HC) |
| 97 | hM4-mCherry | CNO | CNO | CNO | CNO (HC) |
| 102 | hM4-mCherry | CNO | CNO | CNO | CNO (HC) |
| 109 | hM4-mCherry | CNO | CNO | CNO | CNO (HC) |
| 128 | hM4-mCherry | CNO | CNO | CNO | CNO (HC) |
| 113 | PBS | Veh | Veh | Veh | Veh (RS) |
| 116 | PBS | Veh | Veh | Veh | Veh (RS) |
| 137 | PBS | Veh | Veh | Veh | Veh (RS) |
| 101 | hM4-mCherry | Veh | Veh | Veh | Veh (RS) |
| 105 | hM4-mCherry | Veh | Veh | Veh | Veh (RS) |
| 107 | hM4-mCherry | Veh | Veh | Veh | Veh (RS) |
| 115 | PBS | CNO | CNO | CNO | CNO (RS) |
| 118 | PBS | CNO | CNO | CNO | CNO (RS) |
| 120 | PBS | CNO | CNO | CNO | CNO (RS) |
| 123 | PBS | CNO | CNO | CNO | CNO (RS) |
| 124 | PBS | CNO | CNO | CNO | CNO (RS) |
| 131 | PBS | CNO | CNO | CNO | CNO (RS) |
| 135 | PBS | CNO | CNO | CNO | CNO (RS) |
| 94 | hM4-mCherry | CNO | CNO | CNO | CNO (RS) |
| 96 | hM4-mCherry | CNO | CNO | CNO | CNO (RS) |
| 103 | hM4-mCherry | CNO | CNO | CNO | CNO (RS) |
| 108 | hM4-mCherry | CNO | CNO | CNO | CNO (RS) |
| 111 | hM4-mCherry | CNO | CNO | CNO | CNO (RS) |
| 126 | hM4-mCherry | CNO | CNO | CNO | CNO (RS) |
| 129 | hM4-mCherry | CNO | CNO | CNO | CNO (RS) |

**MSPECT-HPLC**

We summarize the elution gradient used in the HPLC for CORT assessment (**Table S**) and the parameters used in the mass spectrometer to ionize, select, fragment and identify CORT and its deuterated derivative (**Table S**).

**Table S2: Elution gradient for corticosterone assessment through HPLC column.**

| Retention (min) | Debit (ml/min) | %phase mobile B |
|---|---|---|
| **0** | 0,150 | 0 |
| **1** | 0,150 | 0 |
| **3** | 0,150 | 25 |
| **10** | 0,150 | 30 |
| **12** | 0,150 | 98 |
| **14** | 0,150 | 98 |
| **15** | 0,150 | 0 |
| **19** | 0,150 | 0 |

**Table S3: Mass spectrometer ionization, selection, fragmentation, and identification parameters.**

| Compound | Polarity | Precursor (m/z) | Product (m/z) | Collision Energy (V) | RF Lens (V) |
|---|---|---|---|---|---|
| **Corticosterone** | positive | 347.11 | 293.472<br>311.294<br>329.169 | 17.03<br>15.91<br>14.95 | 227.49 |
| **D4-corticosterone** | positive | 351.179 | 297.103<br>315.183<br>333.24 | 17.68<br>16.88<br>15.56 | 223.55 |

**Quantification of c-Fos+ cells density**

***Outlining of the targeted areas***

We evaluated c-Fos+ cells density across 56 brain structures along the anteroposterior axis of the brain (see coordinates in **Table S**) and according to the boundaries drawn in the Paxinos and Watson stereotaxic atlas (Paxinos et Watson, 2007), such as shown in **Figure S1** and **Figure S2**.

#### Table S4: Anteroposterior coordinates of the structures investigated

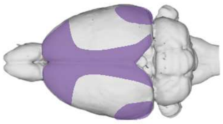

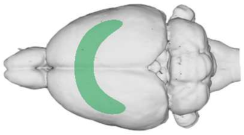

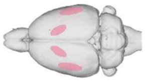

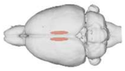

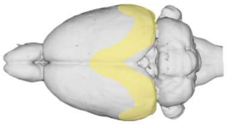

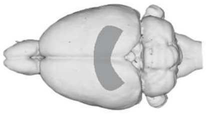

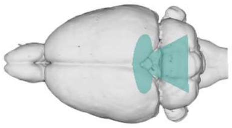

| **Structures** | **Start** *(mm from bregma)* | **End** *(mm from bregma)* |
|---|---|---|
| ACC | 4.20 | 2.52 |
| PRL | 4.68 | 2.52 |
| IL | 4.20 | 2.52 |
| LO | 4.68 | 3.00 |
| MO | 4.68 | 4.20 |
| VO | 4.68 | 3.00 |
| MCC | 2.28 | -1.56 |
| Ins_R | 4.20 | -0.36 |
| Ins_C | -0.36 | -2.92 |
| Cl | 4.20 | -1.08 |
| RS | -1.72 | -6.60 |
| Ect | -3.00 | -6.60 |
| Ent | -3.12 | -6.60 |
| PRh | -3.00 | -6.60 |
| dlS | 2.04 | 0.00 |
| vlS | 2.04 | 0.00 |
| dmS | 2.04 | 0.00 |
| vmS | 2.04 | 0.00 |
| cNac | 2.04 | 0.48 |
| sNac | 2.04 | 0.48 |
| VP | 2.76 | -0.72 |
| LS | 2.28 | -0.60 |
| MS | 1.56 | 0.00 |
| TS | -0.24 | -1.08 |
| lBNST | 0.36 | -0.72 |
| mBNST | 0.72 | -1.08 |
| BA | -1.56 | -3.36 |
| LA | -1.56 | -3.36 |
| CeA | -1.56 | -3.12 |
| MHb | -2.04 | -4.36 |
| LHbL | -2.92 | -4.36 |
| LHbM | -2.92 | -4.36 |
| dCA1 | -2.28 | -6.24 |
| dCA2 | -2.28 | -6.24 |
| dCA3 | -2.28 | -6.24 |
| dDG | -2.28 | -6.24 |
| vCA1 | -4.68 | -6.24 |
| vCA2 | -4.68 | -6.24 |
| vCA3 | -4.68 | -6.24 |
| vDG | -4.68 | -6.24 |
| PVT | -1.20 | -3.72 |
| MD | -1.80 | -3.72 |
| Re | -1.20 | -3.96 |
| LH | -1.56 | -4.56 |
| PVH | -1.56 | -3.36 |
| MM | -4.20 | -5.28 |
| SuMM | -4.20 | -4.80 |
| PAG | -4.56 | -8.52 |
| DR | -6.84 | -9.36 |
| MR | -7.08 | -8.76 |
| SNc | -4.92 | -6.48 |
| SNr | -4.56 | -6.72 |
| LC | -9.48 | -10.08 |
| LDT | -8.16 | -9.36 |
| VTA | -4.68 | -6.48 |
| tVTA | -6.12 | -6.96 |

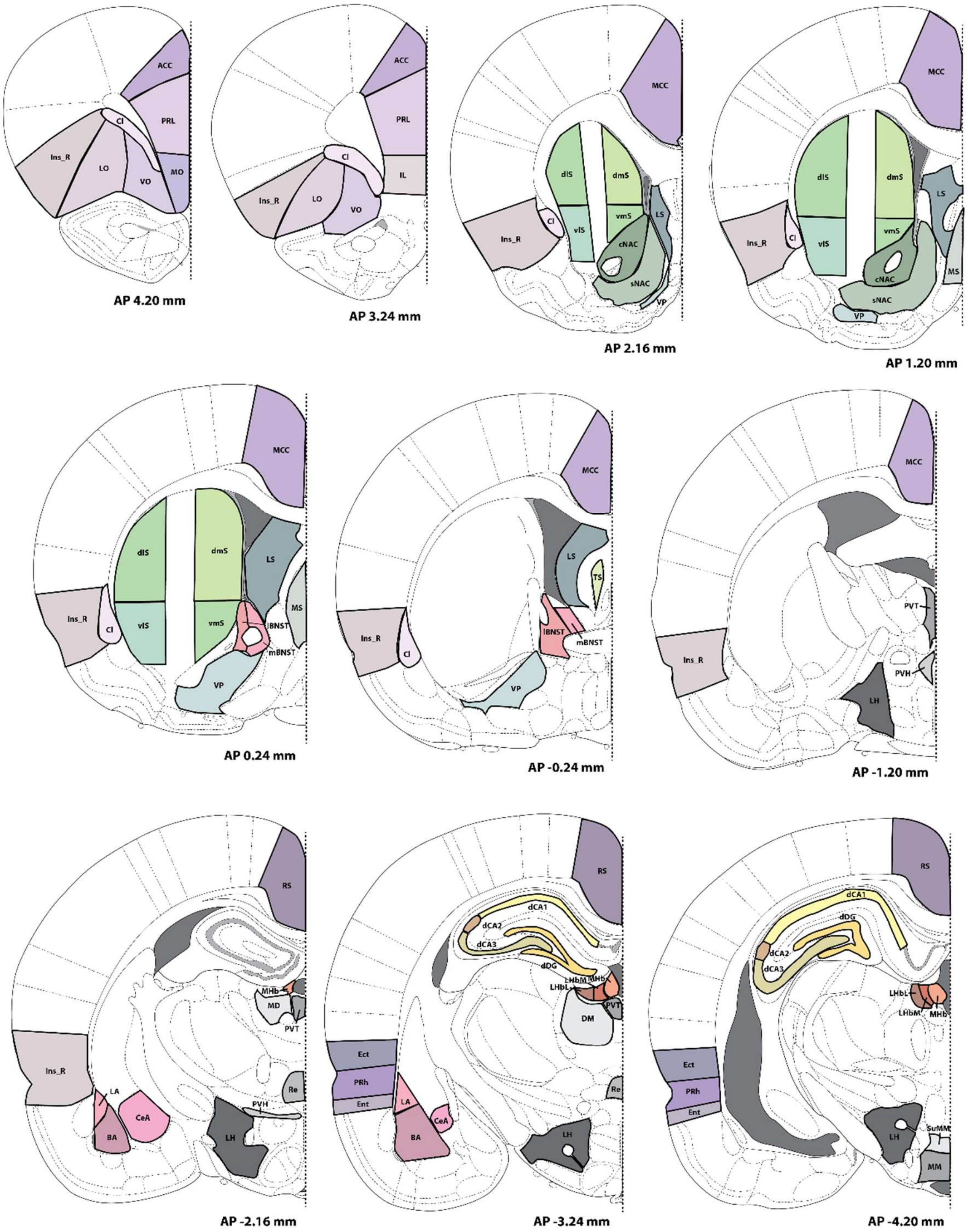


**Figure S1: Boundaries of the structures for the evaluation of c-Fos+ cells densities (part 1)**
Adapted from Paxinos and Watson atlas (Paxinos et Watson, 2007).

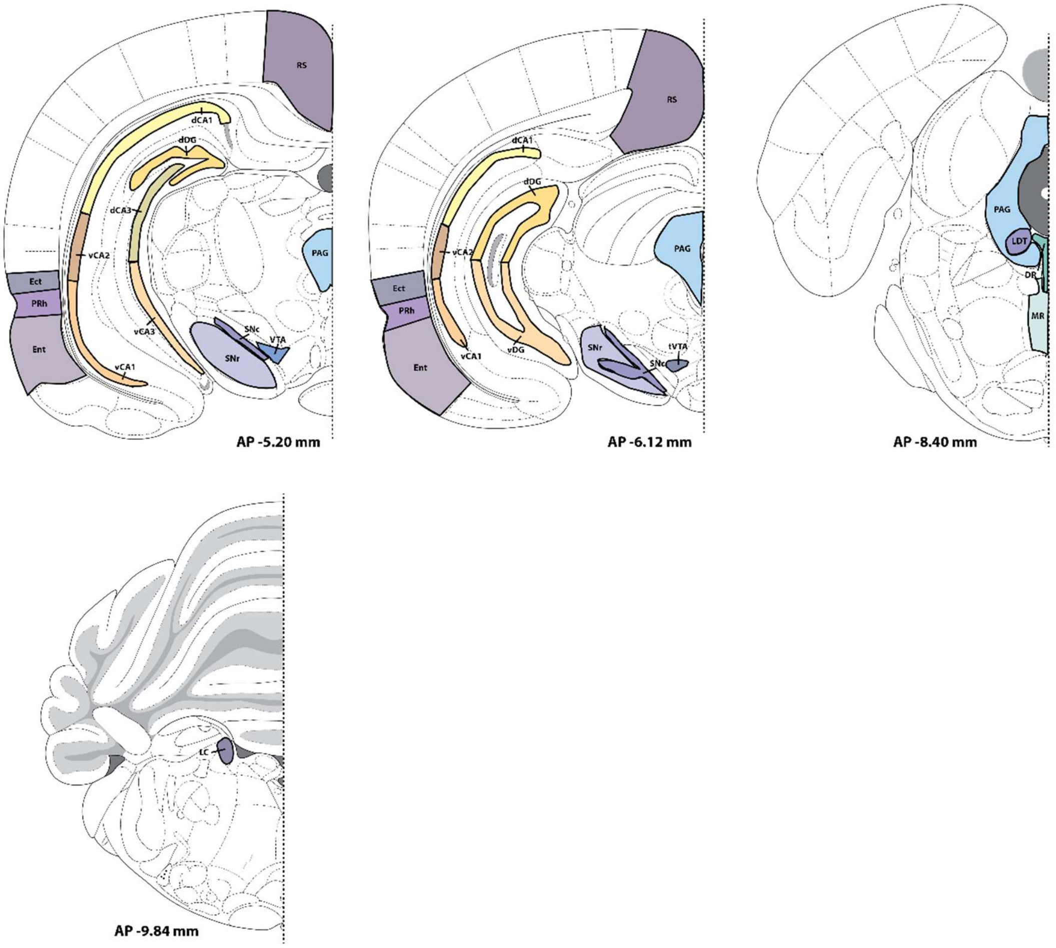


**Figure S2: Boundaries of the structures for the evaluation of c-Fos+ cells densities (part 2)**. Adapted from Paxinos and Watson atlas (Paxinos et Watson, 2007).

***Semi-automated c-Fos+ cells quantification method***

Quantification was performed using ImageJ (Free license, Wayne Rasband, Research Services Branch, NIH, Bethesda, Maryland, USA) with homemade scripts, such as described in Durieux et al., 2020. However, we still verified the accuracy of the semi-automated counting method by comparing it to manual counting, choosing the LHb (**Figure S3**). We found almost identical results with both methods with a correlation $R^2$ = 0.98 (Pearson) and a slope = (1.009 x X) ± 0.0101 (where slope is Y = (a * X) + b [where *Y* is the value given by automated counting, *X* is the value given by manual counting, *a* is the slope of the best fitting line, and *b* is the intersection with the Y axis]. A parameter *a* near 1 suggests that *n* cells counted semi-automatically will be equal to *n* counted manually; a parameter *b* near 0 means that 0 detected cells in the manual counting gives 0 detected cells with the semi-automated method.

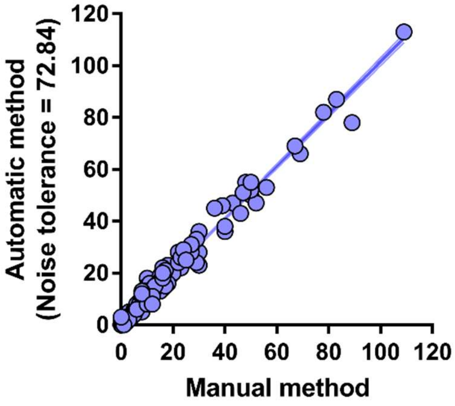


**Figure S3: Assessment of the accuracy of the semi-automated counting method in comparison with the manual counting method.** Plot representing the linear correlation of manual *vs* semi-automated counting; "a" is close to 1, demonstrating the similarity between manual and semi-automated counting, with a very high correlation between both methods ($R^2 = 0.98$; $p < 0.0001$), with the noise tolerance parameter of the FindMaxima() function of ImageJ set to 72.84.

***Resolution of Louvain algorithm***

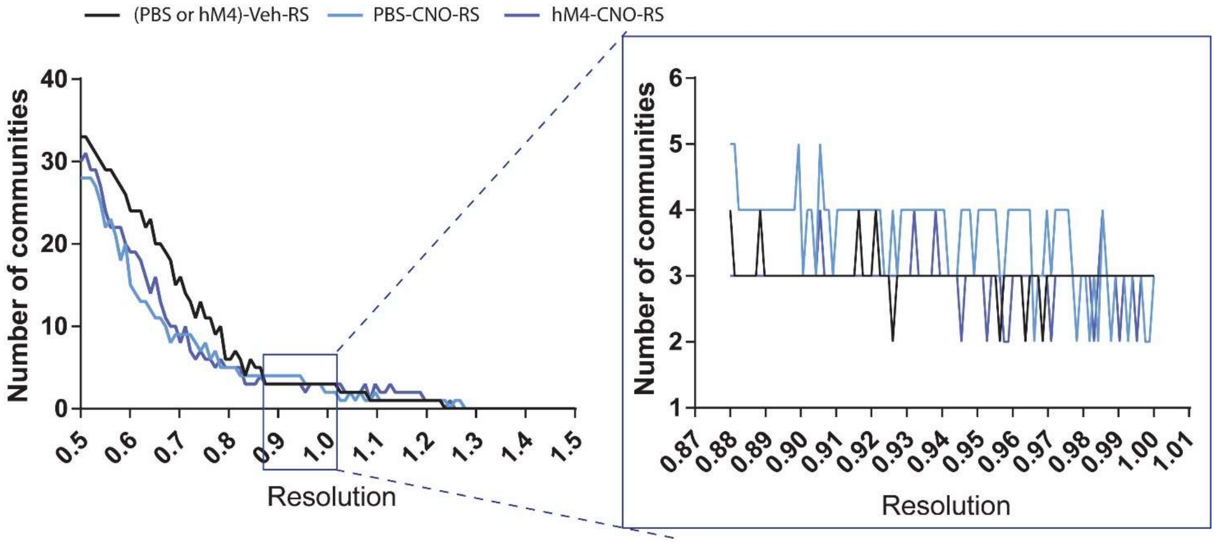


**Figure S4: Plots of the number of communities by the Louvain resolution.** The graphic on the left displays the number of communities extracted in function of the resolution of the Louvain algorithm. We decided to variate the resolution of Louvain Algorithm, staying around the number of communities obtain when the resolution equal 1 (greedier parameter), leading to a variation of the number of communities between 2 and 5. So, we select the interval for the resolution from 0.88 to 1 (as displayed on the right plot). The variation of the number of communities through the resolution interval are displayed by group and over 100 repetitions.

***Within-module strength z-score calculation***

To calculating the strength of the interaction between structures of the same community and between structures of different communities, we evaluated the within-module strength z-score using the following equation:

$$WM_{z-sco} = \frac{Strength_{within} - Strength_{tot}}{std(Strength_{tot})}$$

where $WM_{z-score}$ is the within-module strength z-score for one structure in its own community, $Strength_{withi}$ is the strength (sum of edge weights connecting to the targeted node) considering only the edges connecting with the node (structures) included in the same community than the node targeted, $Strength_{tot}$ is the total strength of the network, and the $std$(x) is the standard deviation of $x$.

### Results

#### Histology

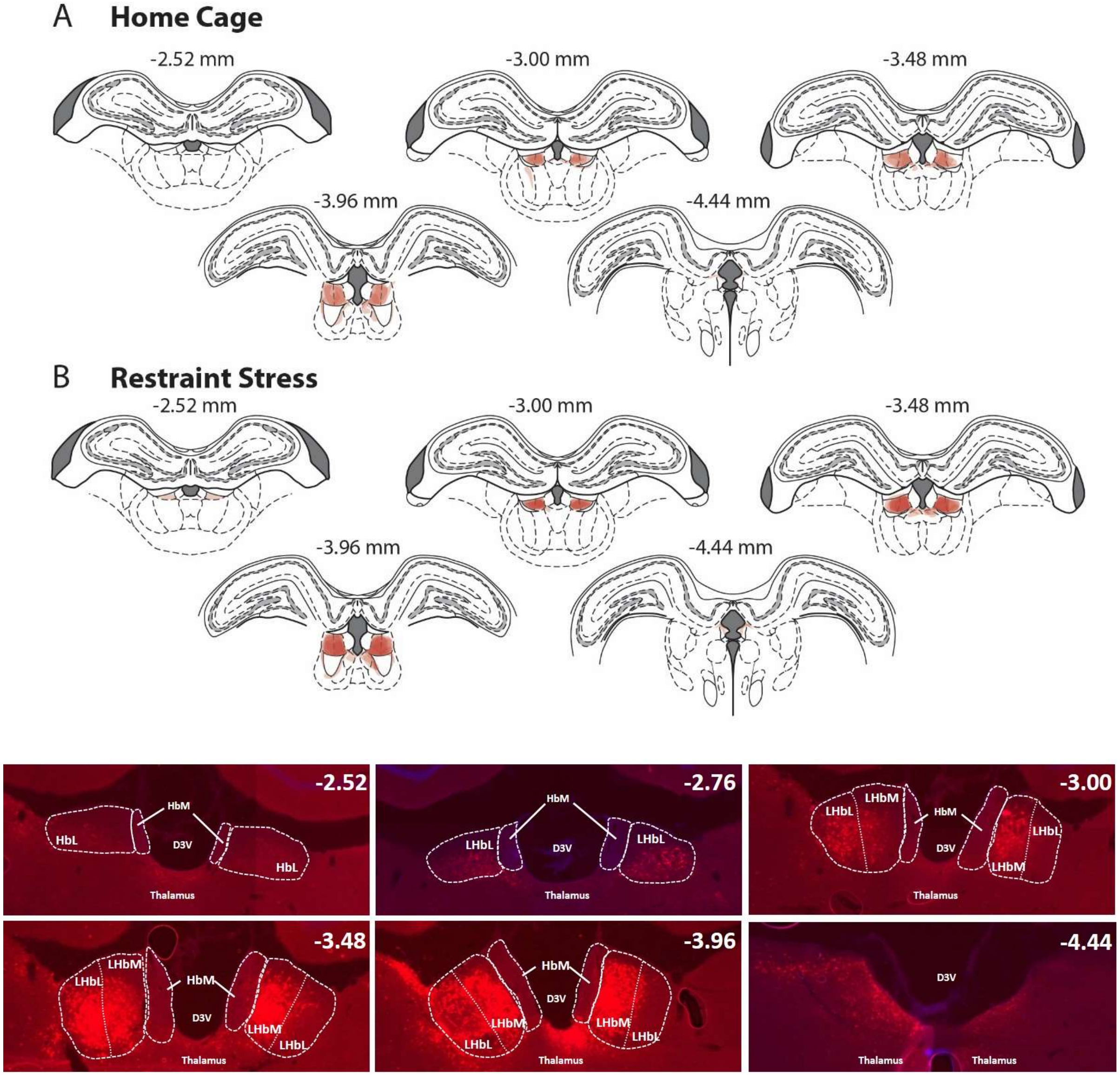


**Figure S5: Schematic representation of the presence of hM4(Gi) receptors in the animals kept following histological verification**. (**Top**) For each rat, on each slide used, the area including the expression of the hM4(Gi) receptors was delineated using a pale red color (opacity 20%). Then, for each stereotaxic coordinate [numbers above the slides correspond to AP coordinates from Bregma (mm) (Paxinos and Watson, 2007)], the slides of all the animals were piled, creating a color scale from pale red to dark red. Therefore, the darker is the area, the greater is the number of animals presenting an expression of the hM4(Gi) receptors within this area. One can see that hM4(Gi) receptors were expressed within the LHb in its entire rostrocaudal extent. The extent of mCherry expression is represented for Home cage rats (**A**; n = 4) and rats subjected to restraint stress (**B**; n = 7). (**Bottom**) Microphotographs of typical example of hM4(Gi) expression within the LHb. White numbers in the top right corner of each photograph are AP coordinates from Bregma (mm). Abbreviations: D3V, third ventricle; LHb, lateral habenula; LHbL, lateral part of the lateral habenula; LHbM, the medial part of the lateral habenula; MHb, medial habenula.

#### C-Fos + cells density raw data and statistics

**Table S5:** Raw c–Fos+ cells density represented by the mean (± SEM) for each region.

| | Home Cage | | | | | | Restraint Stress | | | | | |
|---|---|---|---|---|---|---|---|---|---|---|---|---|
| | (PBS-hM4)-Veh | | PBS-CNO | | hM4-CNO | | (PBS-hM4)-Veh | | PBS-CNO | | hM4-CNO | |
| | *Mean* | *SEM* | *Mean* | *SEM* | *Mean* | *SEM* | *Mean* | *SEM* | *Mean* | *SEM* | *Mean* | *SEM* |
| **ACC** | 0.721 | 0.215 | 1.092 | 0.266 | 10.895 | 4.130 | 37.704 | 8.332 | 27.677 | 3.381 | 53.414 | 14.489 |
| **PRL** | 4.909 | 1.009 | 4.309 | 0.567 | 23.120 | 6.088 | 85.988 | 11.702 | 61.113 | 6.440 | 97.203 | 17.734 |
| **IL** | 13.323 | 2.804 | 15.317 | 3.265 | 34.509 | 9.930 | 92.791 | 10.439 | 67.882 | 6.346 | 93.195 | 14.321 |
| **LO** | 1.280 | 0.275 | 1.220 | 0.268 | 17.253 | 6.337 | 53.287 | 10.291 | 52.767 | 12.463 | 86.572 | 12.218 |
| **MO** | 2.694 | 1.339 | 1.088 | 0.418 | 7.046 | 0.949 | 52.228 | 11.051 | 33.449 | 4.628 | 48.298 | 9.671 |
| **VO** | 1.649 | 0.657 | 1.557 | 0.623 | 19.598 | 8.262 | 68.375 | 11.903 | 63.157 | 13.148 | 98.303 | 14.192 |
| **MCC** | 21.753 | 11.620 | 7.153 | 1.842 | 202.504 | 9.954 | 214.155 | 47.953 | 219.380 | 19.598 | 302.079 | 60.292 |
| **Ins_R** | 1.599 | 0.580 | 1.374 | 0.267 | 8.509 | 3.962 | 28.724 | 2.473 | 25.595 | 4.027 | 41.423 | 3.840 |
| **Ins_C** | 3.860 | 1.492 | 2.048 | 0.474 | 30.214 | 14.465 | 54.409 | 13.275 | 40.551 | 5.877 | 60.889 | 11.558 |
| **Cl** | 16.560 | 3.763 | 7.371 | 1.918 | 74.348 | 26.511 | 125.236 | 28.053 | 102.424 | 13.261 | 135.979 | 14.122 |
| **RS** | 4.345 | 1.687 | 2.492 | 0.600 | 33.303 | 7.722 | 57.801 | 12.095 | 56.221 | 5.425 | 79.778 | 17.365 |
| **Ect** | 4.525 | 0.938 | 3.031 | 0.543 | 13.075 | 3.462 | 59.610 | 6.501 | 43.687 | 6.286 | 54.799 | 12.532 |
| **Ent** | 4.140 | 1.168 | 3.040 | 0.630 | 18.761 | 4.022 | 52.791 | 7.045 | 34.366 | 5.455 | 47.175 | 9.751 |
| **PRh** | 4.256 | 0.731 | 1.914 | 0.289 | 15.420 | 4.056 | 64.873 | 7.885 | 47.579 | 5.515 | 68.570 | 16.136 |
| **dlS** | 0.786 | 0.257 | 0.618 | 0.050 | 2.794 | 1.289 | 1.039 | 0.173 | 1.132 | 0.155 | 3.542 | 1.094 |
| **vlS** | 0.540 | 0.133 | 0.809 | 0.240 | 2.174 | 0.741 | 0.661 | 0.141 | 0.692 | 0.203 | 4.056 | 0.812 |
| **dmS** | 1.617 | 0.471 | 2.238 | 0.562 | 8.224 | 1.222 | 15.559 | 3.533 | 14.059 | 1.922 | 15.943 | 2.208 |
| **vmS** | 1.498 | 0.543 | 1.309 | 0.411 | 6.329 | 1.136 | 12.834 | 2.960 | 10.019 | 1.843 | 13.450 | 1.230 |
| **cNac** | 2.889 | 0.888 | 2.209 | 0.541 | 11.387 | 3.230 | 18.264 | 2.085 | 16.322 | 2.882 | 17.172 | 2.061 |
| **sNac** | 3.065 | 0.769 | 3.724 | 0.769 | 16.255 | 2.694 | 36.828 | 6.436 | 27.652 | 6.506 | 38.231 | 6.772 |
| **PV** | 0.735 | 0.257 | 0.820 | 0.172 | 4.470 | 0.806 | 7.242 | 2.186 | 5.996 | 0.625 | 8.955 | 1.453 |
| **LS** | 5.294 | 1.656 | 3.639 | 0.809 | 28.208 | 7.579 | 67.091 | 10.040 | 45.916 | 4.769 | 62.896 | 8.023 |
| **MS** | 2.371 | 0.780 | 2.469 | 0.601 | 8.091 | 1.071 | 37.899 | 6.623 | 23.527 | 3.478 | 35.374 | 6.399 |
| **TS** | 6.830 | 3.382 | 3.725 | 1.204 | 27.402 | 11.218 | 28.972 | 10.272 | 33.926 | 5.024 | 46.838 | 5.558 |
| **lBNST** | 8.461 | 2.705 | 6.704 | 1.924 | 32.901 | 5.350 | 39.662 | 6.431 | 37.345 | 4.404 | 51.495 | 9.610 |
| **mBNST** | 11.754 | 2.506 | 8.746 | 2.285 | 33.118 | 4.396 | 73.715 | 10.892 | 74.231 | 11.924 | 81.224 | 6.711 |
| **BL** | 5.094 | 1.934 | 2.701 | 0.491 | 25.625 | 8.139 | 44.653 | 8.312 | 37.192 | 5.348 | 45.794 | 7.543 |
| **LA** | 4.272 | 0.954 | 3.278 | 0.562 | 12.311 | 6.145 | 37.839 | 5.710 | 36.060 | 4.238 | 29.892 | 3.907 |
| **CeA** | 7.236 | 2.527 | 18.539 | 4.198 | 36.367 | 7.198 | 45.515 | 10.174 | 45.929 | 3.530 | 83.367 | 20.339 |
| **MHb** | 1.649 | 0.714 | 2.461 | 0.859 | 7.559 | 1.986 | 10.042 | 2.902 | 15.223 | 4.029 | 8.832 | 2.362 |
| **LHbL** | 7.126 | 4.916 | 3.202 | 1.312 | 28.144 | 5.336 | 36.670 | 7.581 | 23.499 | 9.080 | 96.880 | 16.507 |
| **LHbM** | 5.771 | 1.272 | 12.621 | 6.989 | 123.519 | 32.485 | 110.652 | 26.764 | 85.647 | 13.443 | 337.576 | 40.965 |
| **dCA1** | 8.213 | 2.894 | 5.050 | 1.004 | 32.924 | 4.137 | 46.543 | 13.538 | 64.920 | 11.123 | 59.659 | 12.860 |
| **dCA2** | 31.886 | 11.165 | 9.676 | 3.545 | 37.394 | 10.068 | 20.370 | 7.460 | 18.386 | 4.007 | 9.591 | 3.036 |
| **dCA3** | 19.467 | 6.865 | 11.830 | 2.529 | 54.663 | 14.803 | 52.982 | 8.202 | 57.807 | 8.135 | 71.694 | 10.754 |
| **dDG** | 12.399 | 4.181 | 7.978 | 2.078 | 17.048 | 5.345 | 26.870 | 5.155 | 35.962 | 6.915 | 48.545 | 13.084 |
| **vCA1** | 10.078 | 5.444 | 3.423 | 1.023 | 10.944 | 5.513 | 33.651 | 4.641 | 18.746 | 4.350 | 25.917 | 6.289 |
| **vCA2** | 13.167 | 7.693 | 6.871 | 2.124 | 7.822 | 1.536 | 31.464 | 5.859 | 22.900 | 6.848 | 21.833 | 3.464 |
| **vCA3** | 6.590 | 4.001 | 2.788 | 0.947 | 7.236 | 1.921 | 16.642 | 4.575 | 9.298 | 1.981 | 9.326 | 2.169 |
| **vDG** | 2.030 | 0.306 | 2.231 | 0.793 | 3.902 | 2.149 | 11.289 | 3.502 | 7.816 | 2.108 | 9.097 | 2.151 |
| **PVT** | 56.328 | 24.931 | 46.498 | 10.383 | 236.460 | 20.366 | 240.248 | 42.837 | 266.718 | 28.247 | 362.334 | 51.632 |
| **MD** | 0.828 | 0.357 | 1.323 | 0.448 | 53.558 | 26.284 | 10.049 | 1.587 | 5.810 | 1.463 | 47.354 | 20.746 |
| **Re** | 13.697 | 9.106 | 5.959 | 1.382 | 132.411 | 27.645 | 72.389 | 18.300 | 70.241 | 12.864 | 170.591 | 36.268 |
| **LH** | 10.867 | 2.713 | 6.559 | 0.947 | 57.317 | 18.163 | 97.442 | 11.193 | 96.276 | 6.781 | 101.064 | 10.834 |
| **PVH** | 42.686 | 20.326 | 24.312 | 5.963 | 99.025 | 23.669 | 196.286 | 41.287 | 178.912 | 24.710 | 191.432 | 23.541 |
| **MM** | 0.981 | 0.468 | 2.971 | 1.376 | 8.425 | 4.656 | 2.632 | 0.660 | 3.799 | 1.031 | 13.639 | 5.733 |
| **SuMM** | 24.141 | 12.938 | 10.956 | 4.282 | 59.548 | 23.217 | 134.138 | 26.551 | 120.753 | 26.555 | 132.483 | 14.089 |
| **PAG** | 17.023 | 5.953 | 10.609 | 2.976 | 37.055 | 21.584 | 47.888 | 7.051 | 39.294 | 6.844 | 53.248 | 10.458 |
| **RD** | 17.802 | 6.646 | 7.881 | 2.490 | 53.718 | 32.327 | 59.531 | 13.790 | 31.439 | 6.434 | 53.098 | 13.835 |
| **RM** | 7.434 | 2.479 | 15.588 | 10.296 | 75.959 | 33.404 | 32.828 | 9.472 | 25.990 | 5.805 | 54.033 | 12.522 |
| **SNc** | 5.214 | 1.940 | 4.864 | 1.299 | 7.757 | 5.110 | 11.472 | 2.875 | 12.433 | 4.377 | 5.912 | 1.540 |
| **SNr** | 1.213 | 0.227 | 2.693 | 0.819 | 3.278 | 1.585 | 3.917 | 1.149 | 3.621 | 0.971 | 2.714 | 0.495 |
| **LC** | 7.483 | 4.553 | 9.736 | 2.642 | 3.294 | 2.329 | 41.242 | 3.741 | 56.439 | 10.187 | 49.093 | 15.143 |
| **LDT** | 19.470 | 9.423 | 8.766 | 3.598 | 37.490 | 29.355 | 33.553 | 15.022 | 57.374 | 16.710 | 36.697 | 12.307 |
| **VTA** | 14.246 | 6.944 | 4.966 | 1.444 | 21.639 | 11.855 | 40.869 | 11.361 | 22.166 | 4.333 | 27.152 | 3.519 |
| **tVTA** | 5.632 | 1.531 | 2.852 | 0.644 | 11.004 | 2.274 | 7.944 | 1.270 | 9.421 | 2.067 | 15.528 | 3.760 |

**Table S6:** c-Fos+ cells density statistics according to Condition (HC vs RS), Group [(PBS or hM4)-Veh vs PBS-CNO vs hM4-CNO) and Condition x Group interaction. *$p < 0.05$; *$p < 0.01$; ***$p < 0.001$; ****$p < 0.0001$; *****$p < 0.00001$; ns, no significant effect. According to the structures showing significant interactions, the relevant between-group statistics are given in the article.

| | **Condition** | | **Group** | | | | | **Interaction** | |
|---|---|---|---|---|---|---|---|---|---|
| | F value | p | F value | p | **(PBS or hM4)-Veh vs hM4-CNO** | **PBS-CNO vs hM4-CNO** | **(PBS or hM4)-Veh vs PBS-CNO** | F value | p |
| **ACC** | 31.261 | ***** | 2.702 | ns | | | | 0.574 | ns |
| **PRL** | 73.977 | ***** | 3.75 | * | * | ** | ns | 0.865 | ns |
| **IL** | 75.107 | ***** | 3.12 | ns | | | | 1.3 | ns |
| **LO** | 58.453 | ***** | 4.414 | * | *** | ** | ns | 0.554 | ns |
| **MO** | 56.99 | ***** | 1.765 | ns | | | | 0.921 | ns |
| **VO** | 66.259 | ***** | 3.701 | * | ** | ** | ns | 0.345 | ns |
| **MCC** | 33.648 | ***** | 9.124 | ** | *** | *** | ns | 1.352 | ns |
| **Ins_R** | 135.506 | ***** | 8.254 | ** | *** | *** | ns | 1.092 | ns |
| **Ins_C** | 30.892 | ***** | 3.848 | * | * | ** | ns | 0.603 | ns |
| **Cl** | 46.8 | ***** | 5.104 | * | * | ** | ns | 1.053 | ns |
| **RS** | 39.772 | ***** | 4.333 | * | ** | ** | ns | 0.078 | ns |
| **Ect** | 62.651 | ***** | 1.377 | ns | | | | 0.658 | ns |
| **Ent** | 54.113 | ***** | 3.114 | ns | | | | 0.213 | ns |
| **PRh** | 57.232 | ***** | 2.088 | ns | | | | 0.417 | ns |
| **dlS** | 2.309 | ns | 19.147 | ** | *** | ** | ns | 0.073 | ns |
| **vlS** | 2.837 | ns | 17.367 | ***** | *** | *** | ns | 2.663 | ns |
| **dmS** | 46.473 | ***** | 2.154 | ns | | | | 1.123 | ns |
| **vmS** | 47.234 | ***** | 3.45 | * | * | ** | ns | 0.811 | ns |
| **cNac** | 48.672 | ***** | 3.041 | ns | | | | 2.889 | ns |
| **sNac** | 40.182 | ***** | 2.548 | ns | | | | 0.739 | ns |
| **PV** | 32.379 | ***** | 4.357 | * | ** | ** | ns | 0.368 | ns |
| **LS** | 82.743 | ***** | 5.735 | ** | * | *** | * | 2.4 | ns |
| **MS** | 62.965 | ***** | 2.522 | ns | | | | 1.55 | ns |
| **TS** | 22.24 | ***** | 5.559 | ** | ** | ** | ns | 0.435 | ns |
| **lBNST** | 31.063 | ***** | 6.61 | ** | ** | *** | ns | 0.667 | ns |
| **mBNST** | 81.755 | ***** | 2.243 | ns | | | | 0.635 | ns |
| **BA** | 43.401 | ***** | 3.697 | * | * | ** | ns | 1.337 | ns |
| **LA** | 78.97 | ***** | 0.098 | ns | | | | 2.494 | ns |
| **CeA** | 17.92 | *** | 4.902 | * | ** | ** | ns | 0.42 | ns |
| **MHb** | 12.724 | ** | 0.795 | ns | | | | 2.514 | ns |
| **LHbL** | 25.437 | **** | 14.102 | **** | | | | 3.39 | * |
| **LHbM** | 43.003 | ***** | 32.414 | ***** | | | | 4.369 | * |
| **dCA1** | 27.678 | ***** | 1.745 | ns | | | | 1.582 | ns |
| **dCA2** | 3.594 | ns | 2.09 | ns | | | | 3.961 | * |
| **dCA3** | 21.336 | **** | 6.483 | ** | ** | ** | ns | 1.447 | ns |
| **dDG** | 15.854 | *** | 1.551 | ns | | | | 0.679 | ns |
| **vCA1** | 21.16 | **** | 2.975 | ns | | | | 0.518 | ns |
| **vCA2** | 13.527 | *** | 1.298 | ns | | | | 0.073 | ns |
| **vCA3** | 7.155 | * | 2.117 | ns | | | | 0.893 | ns |
| **vDG** | 15.347 | *** | 0.416 | ns | | | | 0.57 | ns |
| **PVT** | 39.285 | ***** | 11.053 | *** | *** | *** | ns | 0.939 | ns |
| **MD** | 0.0068 | ns | 9.459 | *** | *** | *** | ns | 0.204 | ns |
| **Re** | 9.951 | ** | 17.431 | ***** | *** | *** | ns | 0.206 | ns |
| **LH** | 104.963 | ***** | 5.667 | ** | | | | 3.971 | * |
| **PVH** | 42.638 | ***** | 1.52 | ns | | | | 0.93 | ns |
| **MM** | 1.051 | | 4.698 | * | ** | ** | ns | 0.273 | ns |
| **SuMM** | 38.516 | ***** | 1.232 | ns | | | | 0.563 | ns |
| **PAG** | 12.148 | ** | 2.604 | ns | | | | 0.36 | ns |
| **RD** | 4.533 | * | 3.839 | * | ns | ns | * | 1.332 | ns |
| **RM** | 0.205 | ns | 7.752 | ** | ** | ** | ns | 1.686 | ns |
| **SNc** | 2.762 | ns | 0.204 | ns | | | | 1.404 | ns |
| **SNr** | 1.909 | ns | 0.242 | ns | | | | 1.499 | ns |
| **LC** | 33.136 | ***** | 0.592 | ns | | | | 0.333 | ns |
| **LDT** | 3.196 | ns | 0.266 | ns | | | | 1.708 | ns |
| **VTA** | 9.294 | ** | 2.768 | ns | | | | 1.164 | ns |
| **tVTA** | 5.919 | * | 5.703 | ** | ** | ** | ns | 0.494 | ns |

**Table S7**: c-Fos+ cells density statistics including post hoc (Newman-Keuls) comparison of the two control groups, i.e. (PBS or hM4)-Veh-HC and (PBS or hM4)-Veh-RS, according to all structures showing a statistical difference following the ANOVA (**Table S6**). *p<0.05, **p<0.01, ***p<0.001.

| | |
|---|---|
| ACC | * |
| PRL | *** |
| IL | *** |
| LO | ** |
| MO | *** |
| VO | *** |
| MCC | ** |
| Ins_R | *** |
| Ins_C | ** |
| Cl | *** |
| RS | ** |
| Ect | *** |
| Ent | *** |
| PRh | *** |
| dmS | *** |
| vmS | *** |
| cNac | *** |
| sNac | *** |
| PV | ** |
| LS | *** |
| MS | *** |
| Lbnst | ** |
| Mbnst | *** |
| BA | *** |
| LA | *** |
| LHbM | * |
| dCA1 | * |
| dCA3 | ** |
| VCA1 | * |
| vDG | * |
| PVT | ** |
| LH | *** |
| PVH | ** |
| SuMM | ** |
| LC | * |
| VTA | 0.054 |

**Comparison of the structure strength in all the group**

Using the classical bootstrap and the permutation test, we evaluated the significant differences between the three groups [(PBS or hM4)-Veh, PBS-CNO and hM4-CNO] exposed to restraint (Erreur ! Source du renvoi introuvable.**8**). No difference was observed between the PBS-CNO and the hM4-CNO group, suggesting that the effects observed on the functional network in hM4-CNO group may come from CNO effects more than, directly, the inactivation of LHb.

**Table S8:** Significant between-groups strength differences. Represents the three possible comparisons between the groups (columns) for each structure (rows). The cells indicated in red represents when both statistical tests (classical bootstrap and permutation test) were displaying a significant difference (*p* < 0.05), in green (per) when a significant difference only occurred after the permutation test (*p* < 0.05), and in yellow (bs) when a significant difference only occurred after the bootstrap test (*p* < 0.05). Note that only significant interactions detected by both tests have been considered in this study.

| | (PBS or hM4)-Veh vs PBS-CNO | PBS-Veh vs hM4-CNO | PBS-CNO vs hM4-CNO |
|---|---|---|---|
| ACC | p<0.05 | ns | ns |
| PRL | ns | ns | ns |
| IL | ns | ns | ns |
| LO | ns | ns | ns |
| MO | ns | ns | ns |
| VO | ns | ns | ns |
| MCC | ns | ns | ns |
| Ins_R | ns | ns | ns |
| Ins_C | ns | p<0.05 | ns |
| Cl | ns | ns | ns |
| RS | ns | ns | ns |
| Ect | ns | ns | ns |
| Ent | ns | ns | ns |
| PRh | ns | ns | ns |
| dlS | p<0.05 | ns | ns |
| vlS | per | ns | ns |
| dmS | ns | ns | ns |
| vmS | ns | ns | ns |
| cNac | ns | ns | ns |
| sNac | ns | ns | ns |
| PV | ns | ns | ns |
| LS | bs | ns | ns |
| MS | ns | ns | ns |
| TS | p<0.05 | bs | ns |
| lBNST | ns | ns | ns |
| mBNST | bs | p<0.05 | ns |
| BL | ns | ns | ns |
| LA | ns | ns | ns |
| CeA | bs | p<0.05 | ns |
| MHb | ns | ns | ns |
| LHbL | ns | ns | ns |
| LHbM | ns | p<0.05 | ns |
| dCA1 | bs | ns | ns |
| dCA2 | bs | ns | ns |
| dCA3 | ns | ns | ns |
| dDG | ns | ns | ns |
| vCA1 | ns | ns | ns |
| vCA2 | ns | ns | ns |
| vCA3 | ns | ns | ns |
| vDG | ns | ns | ns |
| PVT | p<0.05 | ns | ns |
| MD | ns | ns | ns |
| Re | ns | ns | ns |
| LH | p<0.05 | ns | ns |
| PVH | p<0.05 | bs | ns |
| MM | ns | ns | ns |
| SuMM | ns | ns | ns |
| PAG | ns | ns | ns |
| RD | ns | ns | ns |
| RM | bs | ns | ns |
| SNc | ns | p<0.05 | ns |
| SNr | bs | p<0.05 | ns |
| LC | per | per | ns |
| LDT | bs | p<0.05 | ns |
| VTA | p<0.05 | bs | ns |
| tVTA | ns | ns | ns |

#### Example of random network based on the initial data

The random networks have been created by shuffling the columns and the rows of the correlation of the initial matrices (see **Figure S6** for an example).

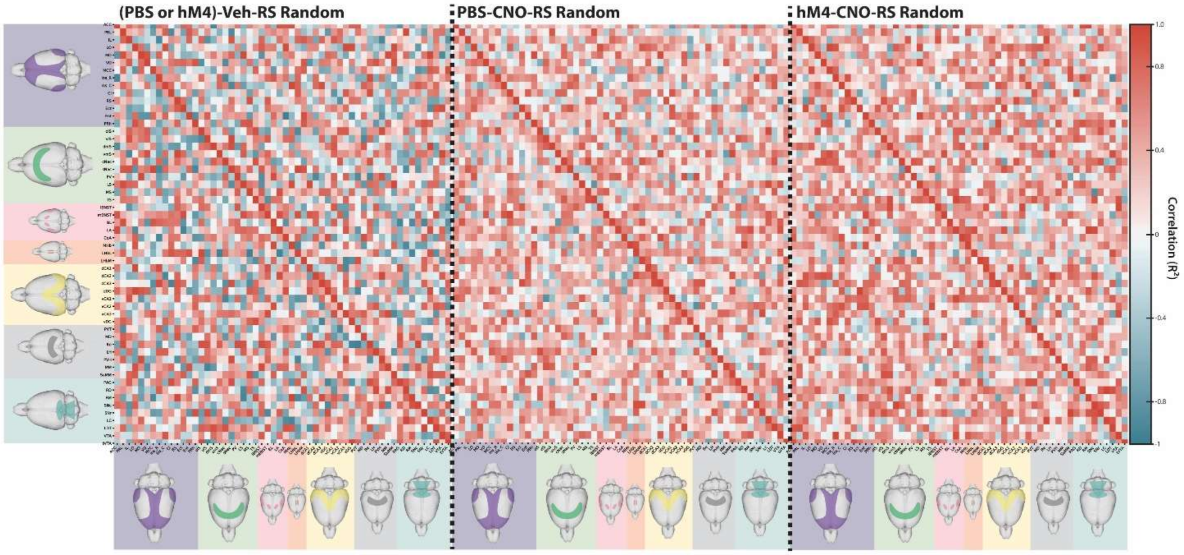


**Figure S6: Example of a random network by group.** Note that 500 random networks have been created for each group to estimate the distribution of the null hypothesis (H0).

#### Global measurement of the network

We observed that the random networks created for each group presented, as planned, the same average clustering and the same average weight (**Figure SA-B**). In addition, they all presented a lower modularity (classic bootstrap $p < 0.05$; **Figure S-C**), suggesting that the networks associated to all groups presented a non-random modularity. This result allowed us to evaluate the allegiance, making the modules accurate and meaningful.

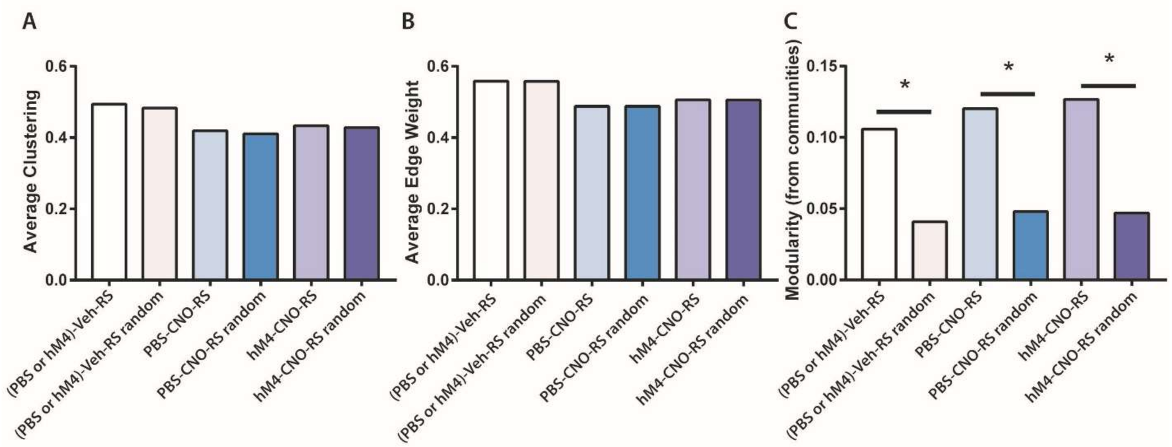


**Figure S7: Global evaluation for random networks associated with the initial groups.** Average clustering (**A**), Average edge weight (**B**) and modularity (**C**) of each group [(PBS or hM4)-Veh-RS, PBS-CNO-RS, hM4-CNO-RS) exposed to restraint and their associated random networks. Statistics: *$p < 0.05$.

**Allegiance analysis on all the groups**

We evaluated the allegiance matrices for each group exposed to restraint (**Figure S**) which served as a base to compare the composition (last main analysis) of their respective communities.

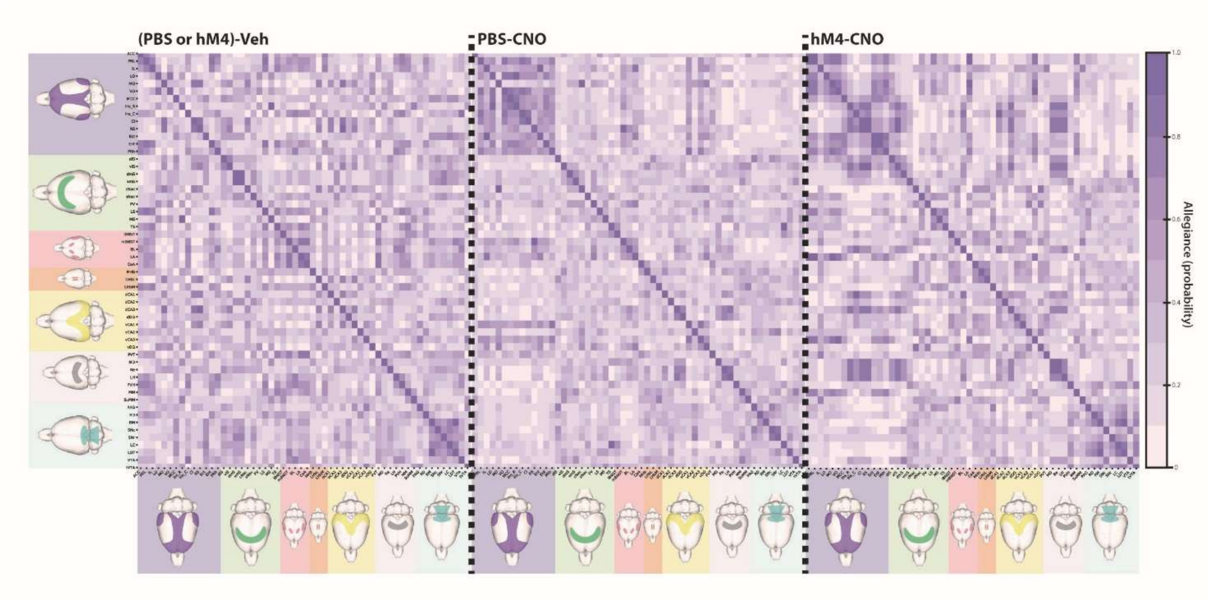


**Figure S8: Allegiance heatmap for the three groups.** Heatmap of allegiance of the (PBS or hM4)-Veh-RS, the PBS-CNO-RS and the hM4-CNO-RS groups, representing the probability for two given structures to belong to the same community across the bootstrap iterations (the darker, the higher probability).

**Compositions of the communities of each group**

**(PBS or hM4)-Veh-RS group:**

Community °1: ACC, PRL, IL, Ins_C, Ent, PRh, LS, TS, lBNST, mBNST, BA, LA, CeA, LHbM, PVT, LH, PVH, MM, DR, VTA.

Community n°2: LO, VO, Ins_R, VP, MS, MHb, dCA2, vCA2, vDG, MD, SuMM, tVTA.

Community n°3: MO, MCC, Cl, RS, Ect, dlS, vlS, dmS, vmS, cNac, sNac, LHbL, dCA1, dCA3, dDG, vCA1, vCA3, Re, PAG, MR, SNc, SNr, LC, LDT.

**PBS-CNO-RS group:**

Community °1: ACC, dlS, vlS, vmS, sNac, LS, TS, lBNST, CeA, MHb, LHbL, LHbM, dCA1, dCA2, vCA2, PVT, MD, PVH, MM, SuMM, PAG, DR, MR, SNc, SNr, LC, LDT, VTA.

Community n°2: PRL, IL, LO, VO, MCC, Ins_R, Ins_C, Cl, RS, Ect, Ent, PRh, mBNST, vCA1, vCA3.

Community n°3: MO, dmS, cNac, VP, MS, BA, LA, dCA3, dDG, vDG, Re, LH, tVTA.

**hM4-CNO-RS group:**

Community °1: ACC, PRL, LO, MO, VO, MCC, Ect, Ent, PRh, sNac, LS, MS, TS, LA, vCA3, MM.

Community n°2: = IL, vmS, cNac, VP, lBNST, mBNST, CeA, MHb, LHbL, LHbM, dDG, vCA1, vCA2, vDG, PVT, PVH, SuMM, PAG, DR, MR, SNr, LC, LDT, tVTA.

Community n°3: Ins_R, Ins_C, Cl, RS, dlS, vlS, dmS, BA, dCA1, dCA2, dCA3, MD, Re, LH, SNc, VTA.
